# 3’UTR cleavage of transcripts localized in axons of sympathetic neurons

**DOI:** 10.1101/170100

**Authors:** Catia Andreassi, Raphaëlle Luisier, Hamish Crerar, Sasja Blokzijl-Franke, Nicholas M. Luscombe, Giovanni Cuda, Marco Gaspari, Antonella Riccio

**Affiliations:** MRC Laboratory for Molecular Cell Biology, University College London, London, WC1E 6BT, UK.; Francis Crick Institute, London, NW1 1AT, UK.; UCL Genetics Institute, University College London, London, WC1E 6BT, UK.; Department of Experimental and Clinical Medicine, University Magna Graecia, Catanzaro, 88100, Italy.

## Abstract

The 3’ untranslated regions (3’UTRs) of messenger RNAs (mRNA) are non-coding sequences that regulate several aspects of mRNA metabolism, including intracellular localisation and translation. Here, we show that in sympathetic neuron axons, the 3’UTRs of many transcripts undergo cleavage, generating both translatable isoforms expressing a shorter 3’UTR, and 3’UTR fragments. 3’end RNA sequencing indicated that 3’UTR cleavage is a potentially widespread event in axons, which is mediated by a protein complex containing the endonuclease Ago2 and the RNA binding protein HuD. Analysis of the *Inositol monophosphatase 1* (*Impa1*) mRNA revealed that a stem loop structure within the 3’UTR is necessary for Ago2 cleavage. Thus, remodeling of the 3’UTR provides an alternative mechanism that simultaneously regulates local protein synthesis and generates a new class of 3’UTR RNAs with yet unknown functions.

---

Messenger RNAs (mRNA) are unique molecules in that they combine a coding sequence carrying the information necessary for the synthesis of proteins, with untranslated regions (UTRs) necessary for transcript localization, translation and stability (1). Many studies have demonstrated that in neurons, mRNA are transported to axons and dendrites where they are rapidly translated into proteins in response to extrinsic stimuli (2). However, despite the extensive analysis of the axonal transcriptome in several neuronal cell types (3, 4), the role of 3’UTR isoforms in regulating mRNA transport and metabolism remains largely unknown.

To examine the distribution of the 3’UTR isoforms of *Inositol monophosphatase 1* (*IMPA1*), a transcript enriched in sympathetic neuron axons (5), we performed 3’ rapid amplification of complementary DNA end (3’ RACE) using mRNA isolated either from cell bodies or distal axons of sympathetic neurons grown in compartmentalized chambers (**Fig. S1A**). In sympathetic neurons, all three IMPA1 isoforms detected expressed an identical coding sequence and 3’UTRs of different length (**Fig. 1A**). Two major isoforms named *IMPA1-Short* (*IMPA1-S*, 3’UTR 1128 nts) and *IMPA1-Long* (*IMPA1-L* contains a 120 nt axonal localization element (5), 3’UTR 1248 nts) were expressed in cell bodies, while the third, newly identified isoform carrying a much shorter 3’UTR was detected only in axons. We named this axon-specific isoform *IMPA1-Cleaved* (*IMPA1-C*, 3’UTR 451 nts, **Fig. 1A**). Northern blot analysis confirmed that the three isoforms were expressed in sympathetic neurons and PC12 cells (**Fig. 1B**). We previously demonstrated that a sequence uniquely found at the 3’end of *IMPA1-L* was necessary and sufficient to localize the transcript to axons (5). To test whether additional elements present in *IMPA1-C* 3’UTR may target the transcript to axons we used a reporter assay based on the expression of a myristoylated, destabilized form of GFP (myrdEGFP) with a very short half-life and limited intracellular diffusion (6, 7). Vectors containing a myrdEGFP coding sequence were flanked by the 3’UTRs of either IMPA1-C (myrdEGFP-IMPA1-C), IMPA1-L (myrdEGFP-IMPA1-L) or as a negative control, histone H3 (myrdEGFP-HH3). As observed previously, when sympathetic neurons were electroporated with myrdGFP-IMPA1-L, the GFP signal was clearly detected in axons up to 1600 μm from the cell bodies. In contrast, the signal from myrdGFP-IMPA1-C was restricted to cell bodies and proximal axons (**Fig. 1C** and **S1B**), indicating that similarly to IMPA1-S (5) and histone H3, the short 3’UTRs lacking the *IMPA1-L* localization element cannot target the transcript to distal axons. Because the 3’ RACE indicated that *IMPA1-C* was expressed in axons, we hypothesized that the short 3’UTR of *IMPA1-C* may be generated by *in situ* cleavage of *IMPA1-L*. Differential expression patterns of isolated 3’UTR fragments and coding sequences has been observed for thousands of neuronal (8, 9) and non-neuronal genes (10, 11). In one instance, small peptides were synthesized from the 3’UTR of genes expressed in aging dopaminergic neurons (12).

**Fig. 1.**
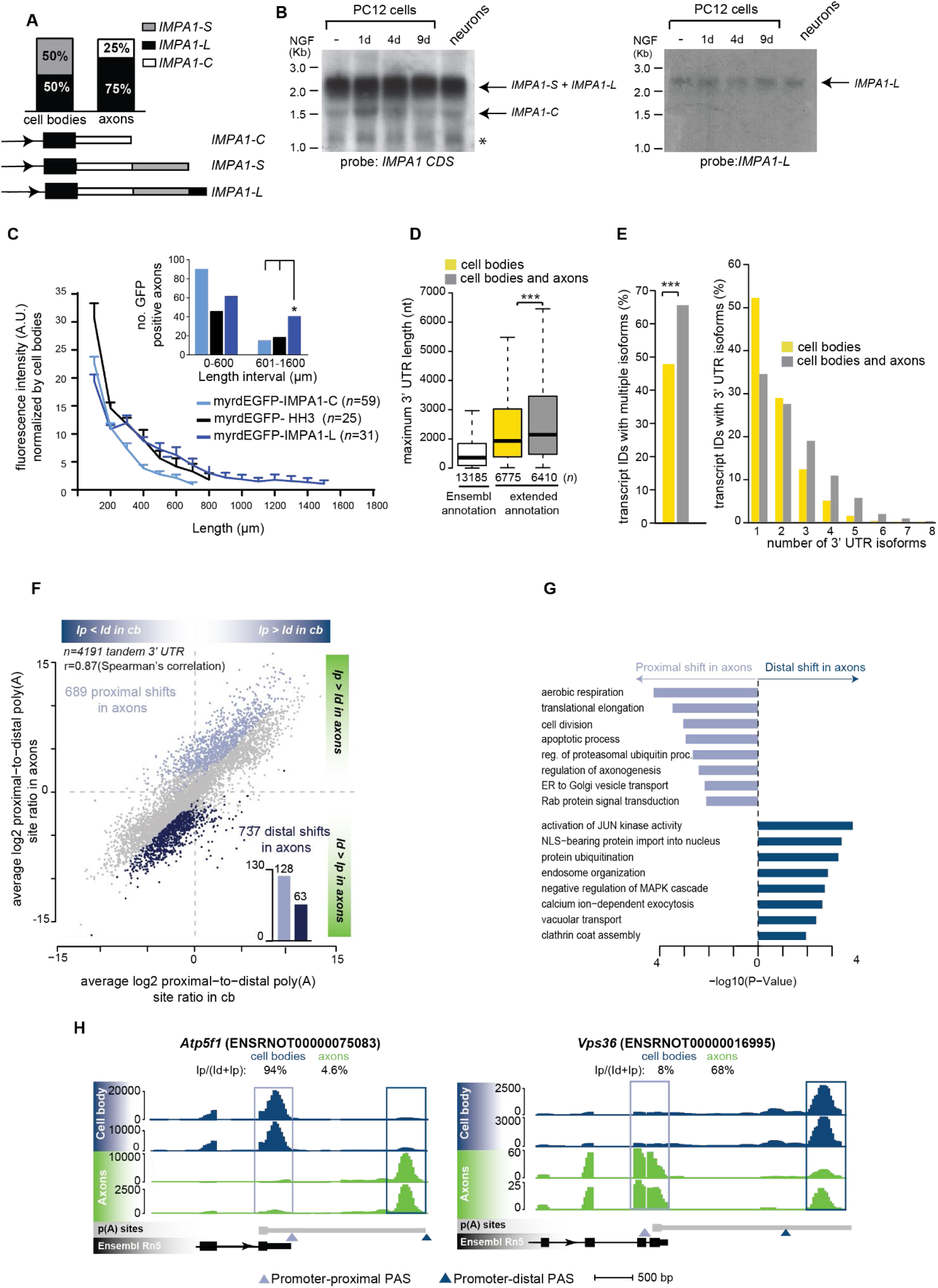
Analysis of 3’UTR PAS choice in axons of sympathetic neurons. (**A**)(*Top*) Percentage of RACE clones containing different IMPA1 3’UTR isoforms from axons and cell bodies as indicated (Cell bodies n=15, axons n=12). (*Bottom*) Schematics of the three IMPA1 3’UTRs. (**B**) Northern blot analysis of RNA isolated from naïve or NGF-differentiated PC12 cells, and sympathetic neurons using 32P-labelled probes annealing with IMPA1 CDS (*Left panel*) or IMPA-L 3’UTR (*Right panel*). Note that the resolution of agarose gels and the abundance of the IMPA1-S isoform prevent the discrimination of IMPA1-S and IMPA1-L when using IMPA1 CDS probe. *= a further isoform with a very short 3’UTR can be detected by IMPA1 CDS probe. (n=3). (**C**) Quantitative analysis of GFP protein immunofluorescence in axons of sympathetic neurons expressing either myrdEGFP-IMPA1-L, myrdEGFP-IMPA1-C, or myrdEGFP-HH3. (Inset): Distribution of GFP-positive axons at the indicated length intervals. (*p<0.05 n=3; Chi-square test). (**D**) Maximum 3’UTR lengths for existing annotations in Ensembl Rn5 and for those newly identified by 3’end RNAseq in this study (*** p < 0.01; Wilcoxon rank sum test). (**E**) Percentage of cell body and axonal transcript IDs showing multiple 3’UTRs (*Left*) and distribution of 3’UTR isoforms per expressed Ensembl transcript ID (*Right*) (*** p<0.01; Fisher exact count test). (**F**) Scatter plot of the relative usage of promoter-proximal and promoter-distal poly(A) sites in cell bodies and axons. (FDR<0.01 between cell body and axonal compartment; Fisher exact test). Dark blue = distal shifts in axons compared to cell body. Light blue = proximal shifts in axons compared to cell body. (Inset) 3’UTR isoforms with proximal or distal shift uniquely detected in axons when we required that usage of promoter-proximal or distal 3’UTR isoform in cell body sample was less than 20% of total isoforms. (**G**) Statistically enriched GO terms for transcripts showing a proximal (*Top*) or distal (*Bottom*) shift in poly(A) site usage in axons. (**H**) Genome browser view of representative transcripts with a marked shift towards decreased (Atp5f1) or increased (Vps36) promoter-proximal poly(A) site usage in axons compared to cell bodies. (See also **Fig. S1–S4**).

To investigate 3’UTR usage in the axonal and cell body transcriptome we performed a stranded 3’end-RNA sequencing on sympathetic neurons. Prior to sequencing, mRNA was subject to two rounds of linear amplification (**Fig. S1C**) that led to the accumulation of reads at the 3’ end of transcripts (**Fig. S2A**) generating a read coverage profile similar to Poly(A)-Seq (13). In addition to the existing Ensembl annotations, we identified 26,468 new 3’UTR isoforms and extended the 3’UTR of 7,506 transcripts (**Fig. S2B-D**). The reliability of these annotations was confirmed by checking them against a comprehensive polyadenylation atlas compiled from a number of independent resources (**Fig. S2E**). Nearly 70% of the newly identified 3’ ends were found within a distance of 100 nt from the annotations in these resources, demonstrating the suitability of our approach (**Fig. S2F**). Analysis of the PAS motifs within 150 nt from the 3’end revealed preferential usage of non-canonical PAS motifs for the longer 3’UTRs (**Fig. S2G**). 3’RACE performed for the *actin beta*, *stathmin 2* and *cofilin1* transcripts on sympathetic neurons indicated that in all cases the isoforms detected matched the 3’ ends identified by the screen (**Fig. S3A, B**). Transcripts were then divided into two categories, those present solely in cell bodies and those also expressed in axons. 9,378 3’UTR isoforms associated with 6,410 transcripts were found in axons (**Fig. S3C-E**). Axonal transcripts expressed longer 3’UTRs and a higher number of 3’UTR isoforms compared to the cell bodies, with many transcripts expressing three or more alternative 3’UTR (**Fig. 1D, E**). Next, we compared the relative usage of promoter-proximal and promoter-distal poly(A) sites (14) between transcripts with multiple isoforms located either in cell bodies or axons. Transcripts containing two or more 3’UTRs isoforms were considered for further analysis (4,191 tandem pairs of 3’UTR isoforms, **Fig. 1F**) and the difference in log2 proximal-to-distal expression ratios of 3’UTR isoforms between cell bodies and axons was calculated. A difference below-1 or above 1, (FDR<0.01, Fisher count test) indicated respectively a distal or proximal shift of poly(A) usage in axons compared with cell body. We found 737 isoforms (17.7% of tandem 3’UTR isoforms) that displayed increased usage of distal 3’UTR isoforms in axons (**Fig. 1F**, dark blue dots) and 689 3’UTR (16.5% of tandem isoforms) that preferentially expressed short isoforms in axons (**Fig. 1F**, light blue dots), with high correlation between sample types (Spearman coefficients r=0.97 for cell bodies samples and r=0.64 for axon samples) (**Fig. S3F**). GO functional analysis revealed that terms associated with axon growth and energy and protein metabolisms were statistically overrepresented among axonal transcripts with shorter 3’UTRs, whereas terms associated with more general biological pathways, such as intracellular signalling, were enriched among axonal transcripts with longer 3’UTRs (**Fig. 1G**). A subset of transcripts selected by applying a thresholding method (see **Fig.1F** legend and Experimental procedures for details) displayed extreme differences in isoform usage, with 63 transcripts with longer 3’UTR and 128 with shorter 3’UTR either uniquely detected or very highly expressed in axons (**Fig. 1F**, inset). Examples of transcripts with strikingly distinct poly(A) usage in cell bodies or axons are shown in **Fig. 1H** and **Fig. S4 A, B**.

Lack of detection of short isoforms in cell bodies may be due to the fact that they are generated co-transcriptionally by alternative polyadenylation and rapidly transported to axons. However, *IMPA1* 3’ RACE (**Fig. 1A**) and the transport assay (**Fig. 1C**) indicate that at least in some cases, isoforms using a short 3’UTRs are expressed in axons but cannot be transported distally, and must be generated in situ by alternative mechanisms, such as cleavage and shortening of long 3’UTRs. To explore this hypothesis, 3’UTR fragments potentially generated by cleavage were detected using RML RT-PCR, a technique that couples 5’P-dependent RNA oligo-Mediated Ligation to RT-PCR followed by cloning of the cleaved fragments (**Fig. S5A**). We reasoned that the predicted cleavage site would likely be in proximity of the proximal PAS as the cleavage would generate an isoform expressing a shorter 3’UTR. RML RT-PCR of mRNA isolated from severed axons (**Fig. S5B**) revealed that when *IMPA1-L*, *Sms* and *Maoa* were tested, most clones contained fragments corresponding to the cleaved 3’UTRs (**Fig. 2A** and **S5C-D**). Remarkably, the fragments were stable, homogenous in size and mapped to precise positions relative to our predicted cleavage site, suggesting that they are not generated by 5’-3’ exonucleolytic degradation. By contrast, 3’UTR cleavage was not detected in transcripts that did not show alternative poly(A) site usage, such as *Cops3*, *Fdxr* and *Maf1* (**Fig. 2B** and **Fig. S5E**). Thus, our findings strongly suggest that most, if not all, axonal specific short isoforms of I*MPA1*, *Maoa* and *Sms* are generated through a process of remodeling the 3’UTR that takes place in axons.

**Fig. 2.**
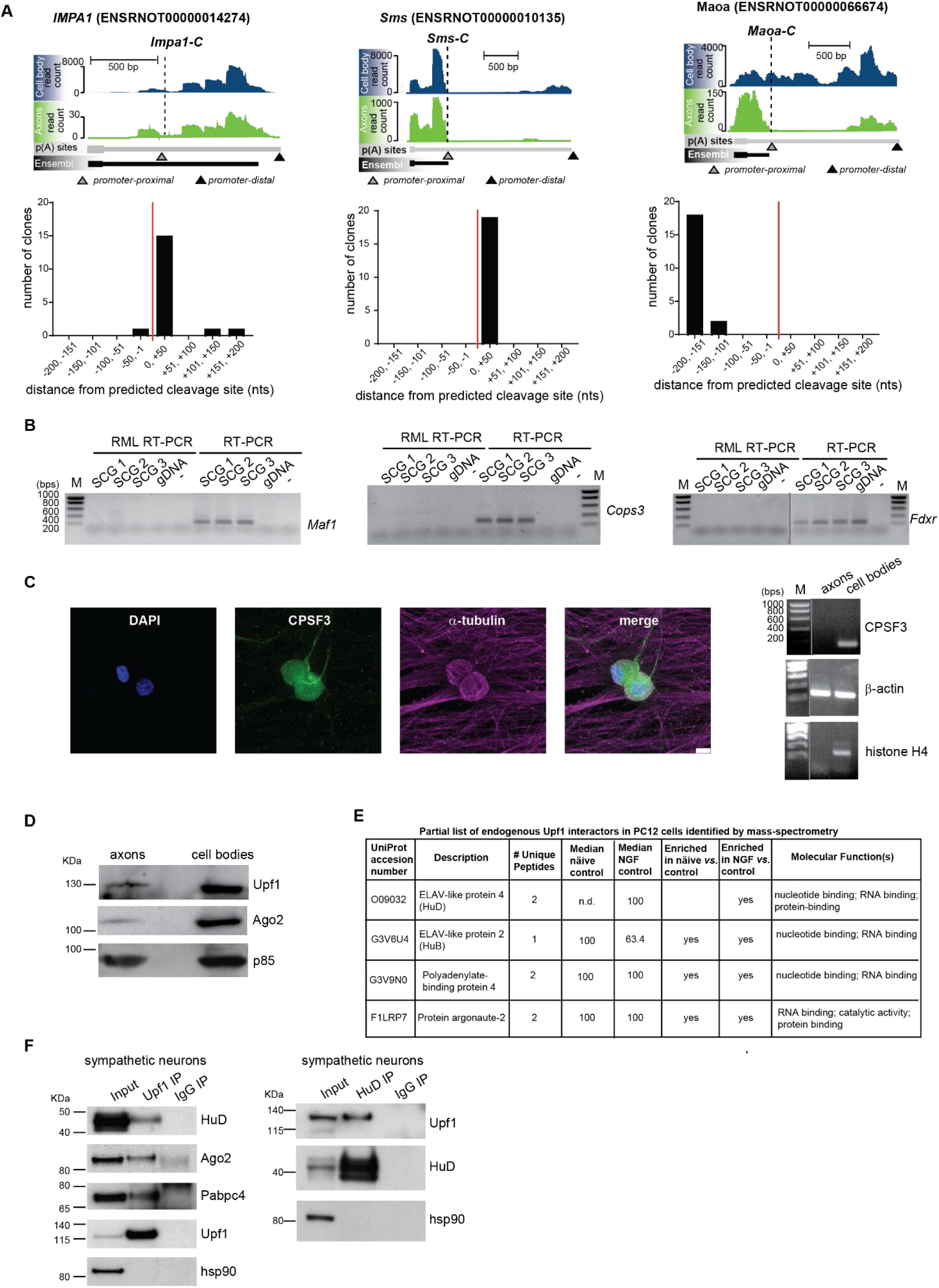
The 3’UTRs of multiple transcripts are remodelled in axons of sympathetic neurons. (**A**) Genome browser view of IMPA1, Sms and Maoa transcripts in axons and cell bodies transcriptomes by 3’end RNA-seq (grey arrowhead). Ensembl annotation (black arrowhead) is also shown. Dashed lines indicate the short 3’UTR isoforms. (*Bottom*) Number of clones of cleaved IMPA1-L, Sms and Maoa 3’UTR fragments purified from axonal RNA grouped according to distance from the predicted cleavage site (red line). Each bin=50nts. (**B**) Absence of cleaved fragments in Maf1, Cops3 and Fdxr transcripts (*left lanes*). The presence of corresponding cDNAs was assessed by regular RT-PCR (*right lanes*). Grey vertical line indicates samples ran on separated gel. (**C**) (*Left*). Representative images of CPSF3 immunostaining in axons and cell bodies of sympathetic neurons. α-tubulin immunostaining was used as a control. Scale bar = 10μm. (*Right*) mRNA isolated from axonal and cell body compartments of sympathetic neurons cultured in compartmentalised chambers was purified and subjected to RT-PCR analysis. The absence of cell body material in axonal samples was assessed using primers amplifying Histone H4 transcripts. (**D**) Western blotting of Upf1, Ago2 and PI3K subunit p85 (as loading control) on axons and cell bodies of sympathetic neurons (n=3). (**E**) Partial list of Upf1 interactors identified in PC12 cells by mass-spectrometry n.d.= not detected. (**F**) Co-immuno-precipitation of Upf1 (*Left*) or HuD (*Right*) with the indicated proteins in sympathetic neurons. (n=3). (See also **Fig. S5** and **S6A-B**).

We next sought to identify the RNA binding proteins that mediate 3’UTR remodeling. A suitable candidate was the nuclear cleavage and polyadenylation specificity factor CPSF3 (15). However, neither *CSPF3* mRNA nor protein were detected in axons (**Fig. 2C**). Mass spectrometry analysis of RBPs associated with polyadenylated transcripts in sympathetic neurons previously performed in our laboratory (Aniko Ludanyi, M.G and A.R,) had revealed that the DNA/RNA helicase Upf1 was one of the few RBPs that interacted with axonal transcripts in response to nerve growth factor (NGF). Upf1 is part of the complex that mediates nonsense-mediated decay (NMD) of mRNA, an RNA surveillance pathway that prevents the translation of truncated proteins by inducing rapid degradation of transcripts harboring a premature termination codon (16). Binding of Upf1 is enriched on longer 3’UTRs and contributes to maintaining mRNAs in a translationally silent state (17). Western blotting of proteins extracts obtained from cell bodies or axons showed that Upf1 is detected in both cellular compartments (**Fig. 2D**). Because RBPs are usually found within large multi-protein complexes, we performed mass spectrometry analysis of proteins that co-immunoprecipitated with Upf1 (**Fig. S6A**). 325 unique peptides that mapped on 72 proteins were identified (Tables S1, S2 and S3), including known interactors of Upf1, such as Polyadenylate-binding protein 1 (Pabp1), ELAV-like protein 2 and interleukin enhancer-binding factor 2 (https://thebiogrid.org/111908/summary/homo-sapiens/upf1.html). Interestingly argonaute-2 (Ago2) and the neuron-specific ELAV-like protein 4 (HuD) were among the most abundant proteins that co-immunoprecipitated with Upf1 (**Fig. 2E**). Co-immunoprecipitation experiments confirmed the interaction of Upf1 with HuD, Ago2 and the PolyA-binding protein cytoplasmic 4 (Pabpc4) in sympathetic neurons (**Fig. 2F**) and PC12 cells (**Fig. S6B**). RNA immunoprecipitation (RIP) assays performed on sympathetic neurons showed a robust interaction of Ago2, HuD and Upf1 with *IMPA1-L* 3’UTR (**Fig. 3A** and **S6C, D**). We detected significantly less binding of Ago2 to *Maf1* 3’UTR (**Fig.3A**), a transcript that is not predicted to undergo 3’UTR cleavage (**Fig. 2B** and **S5E**). Ago2 is the only member of the Argonaute family of proteins with endonuclease activity and binds preferentially to long 3’UTRs (18). To investigate whether Ago2 cleaved *IMPA1* 3’UTR, we performed an *in vitro* cleavage assay. Recombinant Ago2 was incubated with a 5’ end-labelled RNA oligonucleotide encompassing the predicted cleavage site, and sympathetic neuron cell lysate. A stable fragment of the expected size was detected (64 nts, **Fig. 3B** and **S6E**), together with a smaller fragment probably generated by the trimming of the primary cleaved fragment (19). Mutations of the cleavage site identified with the RML RT-PCR assay completely abolished Ago2-dependent cleavage (**Fig. 3B**). Ago2 is a double strand (ds) RNA endonuclease that typically cuts through miRNA paired to target mRNA (20). However, Ago2 can also cleave miRNA precursors and mimetics by recognizing stem-loop structures (21, 22). When *IMPA1-L* 3’UTR was run through the RNA folding prediction software RNAfold, the sequence surrounding the predicted cleavage site formed a stable stem-loop (**Fig. 3C**, WT). To test whether Ago2-dependent cleavage of *IMPA1-L* 3’UTRs may occur through the binding and cleavage of a dsRNA structure, we synthesized oligos with either an impaired stem structure (Δstem) or an enlarged loop (mutant loop). Ago2-dependent cleavage of these mutants was virtually undetectable (**Fig. 3B**), confirming that the stem-loop structure surrounding the cleavage site is necessary for Ago2 cleavage activity. In the Δstem mutant oligo the cleavage site sequence is intact, further indicating that Ago2 cleavage of *IMPA1-L* is independent of miRNA potentially targeting the cleavage site. It should be noted that the cleavage assay was performed in the presence of neuronal cytoplasmic lysates, implying that additional RNA binding proteins and/or co-factors may be necessary either for the cleavage reaction and/or the stabilization of the cleaved fragments. Moreover, 5’RML assay followed by RT-qPCR showed that silencing of Ago2, HuD/B and Pabpc4 decreased the cleavage of endogenous *IMPA1-L* 3’UTR consistent with the levels of siRNA-mediated silencing for each molecule (**Fig. 3C** and **S5F**).

**Fig. 3.**
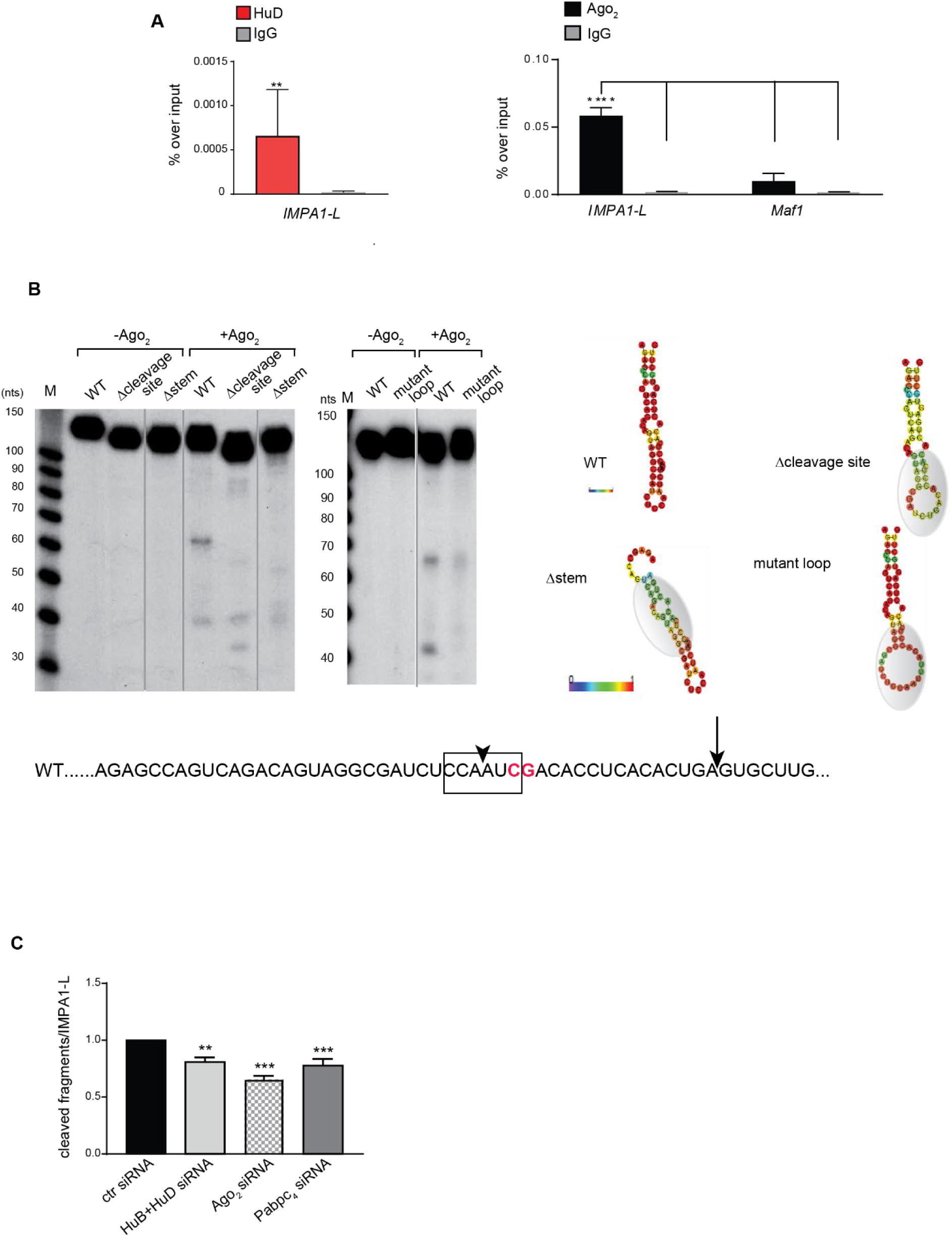
A complex containing Ago2, HuD and Upf1 mediates the cleavage of IMPA1 3’UTR. (**A**) RNA Immunoprecipitation (RIP) of IMPA1-L or Maf1 mRNA with HuD (*Left*) or Ago2 antibody (*Right*), and normal IgG antibody in sympathetic neuron lysate. Data expressed as averages ±s.e.m. (*p<0.05 paired two-tailed t test, ****p<0.0001 One-way ANOVA Dunnett’s multiple comparison test, n=3). (**B**) *In vitro* cleavage assay of radioactive wild type or mutant IMPA1-L 5’ end-labelled RNA oligonucleotides using human recombinant Ago2 and cytoplasmic lysates of sympathetic neurons (*Top Left*). A band corresponding to the expected size of the cleaved fragment (67 nts) is detected in wild type oligonucleotides and is absent when the cleavage site (Δcleavage site) or the secondary structure (Δstem and mutant loop) of the oligonucleotide are mutated. (*Top Right*) Folding predictions of wild type and mutant IMPA1-L oligos. Color-coded probability of pairing is shown. Shadowed area points to the effect of the mutation on the secondary structure of the oligo. (*Bottom*) Sequence of the wild-type oligo used for folding prediction. Arrowhead indicates the point of cleavage. Boxed sequence is deleted in the Δcleavage site mutant. Nucleotides in bold are mutated in the mutant loop oligo. Arrows indicates truncation of the Δstem mutant. (**C**) IMPA1-L cleavage was assayed by RML RT-qPCR on RNA purified from PC12 cells transfected with the indicated siRNAs. Data expressed as averages ± s.e.m. (*** p<0.0005, **p<0.005; one-way ANOVA, Dunnett’s multiple comparison test, n>3). (See also **Fig. S6C-F**).

Given that the 3’UTR length correlates with translation levels in many cell types (23,24), including neurons (26, 27), we reasoned that 3’UTR cleavage could regulate protein synthesis in axons. The isoform generated by the shortening of the 3’UTR (*IMPA1-C*) was polyadenylated as efficiently as *IMPA-L* (**Fig. S7A**) and luciferase assays demonstrated that the 3’UTR of *IMPA1-C* promoted translation at levels similar to *IMPA1-L* (**Fig. 4A**). Mutation of the proximal PAS sites of IMPA1-L (firefly-IMPA1-LΔ poly(A)) inhibited the remodeling of *IMPA1-L* 3’UTR into *IMPA1-C* (**Fig. 4B**) and decreased translation (**Fig.4A**) without affecting mRNA stability (**Fig. S7B**). Polysomal fractionation confirmed a substantial shift of *firefly-IMPA1-LΔ poly(A)* toward the lighter, monosome-rich fractions that is normally associated with lower levels of translation, whereas *firefly-IMPA1-L* and *firefly-IMPA1-C* preferentially co-sedimented with the polysome-enriched fractions (**Fig. S7C** and **Fig. 4C**).

**Fig. 4.**
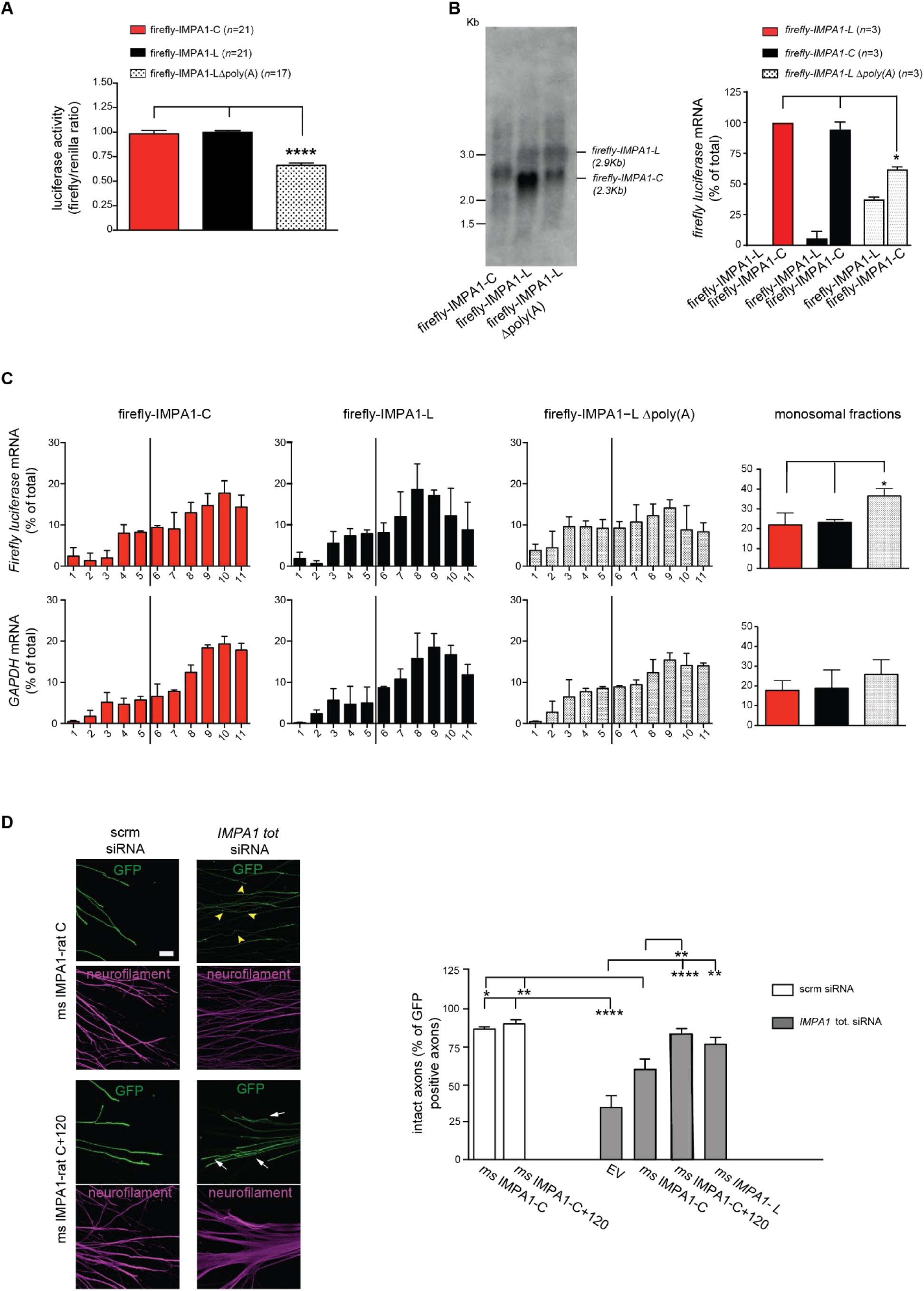
The cleavage of IMPA1 increases the translational efficiency of the transcript. (**A**) PC12 cells were transfected with the indicated Firefly Luciferase vectors and Renilla Luciferase, and subjected to luciferase assay. Data presented as averages ±s.e.m (****p<0.0001; one-way ANOVA, Tukey’s post hoc test, at least 4 independent experiments). (**B**) Firefly luciferase northern blotting of PC12 cells transfected with either Firefly-IMPA1-C, Firefly-IMPA1-L, or Firefly-IMPA1-L Δpoly(A) expression vectors (*Left*). The slight difference in eletrophoretic mobility observed for the firefly-IMPA1-C isoforms (compare lanes 2 to lane 1 and 3) may be due heterogeneity in the poly(A) tail length. (*Right*) Quantitative analysis of Firefly luciferase levels. Data presented as average ±s.e.m. (*p<0.05; multiple t test, using the Holm-Sidak method with α=0.05,. n=3) (**C**) Lysates of PC12 cells transfected with the indicated Firefly-IMPA1 3’UTR vectors were separated by polysomal fractionation, RNA was isolated from each fraction and subjected to northern analysis using 32P-labeled probes to detect Firefly (Upper) or GAPDH (Lower) transcripts. Bands were captured using a phosphorimager and quantified using ImageQuant 5.2. Vertical black bar indicates separation between monosomal and polysomal fractions. (*Right graphs*) Amount of mRNA in cumulative monosomal fractions for lysates transfected with indicated vectors. Data presented as averages ±s.e.m. (*p< 0.01; one-way ANOVA, Tukey’s post hoc test’ n= 3-5). (**D**) (*Left*) Representative images of SCG explants electroporated with scrambled (scrm) siRNA or total IMPA1 siRNA, in the presence of ms HA-IMPA1-C or ms HA-IMPA1-C+120, and GFP. Arrows point to healthy, intact axon bundles, arrowheads to degenerating axon bundles with characteristic beads-on-string appearance. Scale bar 75 μm. (*Right*) Quantitative analysis of the data, presented as average ± s.e.m. (*p<0.05, **p<0.005, ***p≤0.001 and ****p<0.0001; one-way ANOVA, Tukey’s multi-comparison test, n=4). (See also **Fig. S7B-E**).

We previously showed that local synthesis of IMPA1-L in axons is necessary to maintain axon integrity (5). To investigate whether *IMPA1-C* was sufficient to rescue axon degeneration induced by *IMPA1* silencing, we generated rescue vectors containing HA-tagged mouse IMPA1 (*ms* HA-IMPA1) flanked by either IMPA1-L or IMPA1-C 3’UTR, and co-transfected them with siRNA targeting the coding region of rat *IMPA1* (*IMPA1 tot* siRNA, **Fig. S7D**). As previously observed, transfection of *ms* HA-IMPA1-L rescued axon degeneration induced by *IMPA1* silencing (**Fig. 4D** and **Fig. S7E**). Conversely, *ms* HA-IMPA1-C, which cannot be transported to axons did not rescue axon degeneration induced by IMPA1 silencing. When the 120 nts localization sequence was added to IMPA1-C (*ms* HA-IMPA1-C+120) (**Fig. 4D** and **Fig. S7E**) we observed a remarkable increase of axonal survival in neurons lacking IMPA1. These findings confirm that *IMPA1-C* 3’UTR lacks the localization element necessary for axonal transport. They also show that when forcibly targeted to axons, *IMPA1-C* is as efficient as *IMPA1-L* in promoting axon integrity.

Generation of mRNA isoforms bearing alternative 3’UTRs is thought to occur exclusively in the nucleus, where transcriptional elongation is coupled with 5’ end capping, splicing of pre-mRNA, cleavage and polyadenylation of the 3’ end (28). Here, we show that at least for some transcripts, a remodeling of the 3’UTR takes place outside the nucleus and at the site of protein synthesis (**Fig. S8**). While the open reading frame carries the genetic information from the DNA to the translational machinery, cleavage of the 3’UTR generates a class of RNA fragments with yet unknown functions. An implication of our findings is that mRNA transcripts may simultaneously have coding-dependent and coding-independent functions, adding a remarkable layer of complexity to the regulation of gene expression.

## Materials and Methods

### Reagents

Cell culture reagents, molecular biology reagents and kits were purchased from Thermo Fisher Scientific and all other chemicals from Sigma, unless stated otherwise.

### Compartmentalized cultures of sympathetic neurons and SCG explants

All animal studies were approved by the Institutional Animal Care and Use Committees at University College London. Superior cervical ganglia were dissected from post-natal day 1 (P1) Sprague Dawley rats and used for explants or enzymatically dissociated and plated in dishes or in compartmentalized chambers, as previously described (5). SCG explants were cultured on poly(D)lysine-laminin for 9-10 days before surgical removal of cell bodies.

PC12 cells (purchased from ATCC) were maintained in DMEM containing 10% FBS, 5% HS (Hyclone), 2mM glutamine. To induce cell differentiation serum concentration was reduced to 0.5% FBS and 0.25% HS and NGF (50ng/mL) was added for the indicated time. Cells were transfected with Lipofectamine2000 in OptiMEM according to the manufacturer’s instructions.

### RNA isolation, reverse transcription, linear amplification and 3’end-RNASeq

To ensure that the axons were free of cell bodies, prior to each experiment axon compartments were incubated with Hoechst 33342 (10μg/mL in PBS for 20 min at 37°C) and observed under an inverted fluorescent microscope. Cultures showing cell nuclei in the axon compartments or leakage of the dye in the central compartment were discarded. Total axonal and cell bodies RNA was purified from the lateral compartments of 52 or 36 chambers and the central compartment of 7 or 6 chambers respectively obtained from 3 or more independent cultures. Total RNA was isolated using PureLink^®^ RNA Micro Scale Kit, according to the manufacturer’s instructions with minor modifications. Briefly, axons and cell bodies were collected from chambers using lysis buffer (300μl) containing 10% β−mercaptoethanol. Total mRNA bound to the columns was washed and eluted twice in elution buffer (12μl). Aliquots of each sample was reverse transcribed in a 20μL reaction volume containing random hexamer mix and 50U SuperScript III Reverse Transcriptase at 50°C for 1 hr. To check the quality of samples and the absence of cell bodies contamination in axon samples, first-strand cDNAs (5μL) were PCR amplified in a 25μL PCR reaction containing actin beta or histone H4 specific primers (0.20μM), dNTPs (200nM) and Go Taq polymerase (1.25U, Promega). Primer sequences and PCR conditions are provided in Table S4.

For mRNA linear amplification, samples were purified as described above, concentrated by speed-vacuum centrifugation to 1μL (axons) or 5μL (cell bodies) volume, and used for two rounds of linear amplification as previously described (29). The volume of the first-strand reaction for the axons was scaled down to 5μL. After the second round of amplification contaminant cDNA was digested by treating the samples with RNAse-free DNAse (2U, Epicentre). Performance of the samples was tested by RT-PCR. Linear amplified aRNA from cell bodies and axon samples (2 biological replicates each) was used to prepare RNASeq libraries using the strand-specific ScriptSeq protocol (Illumina). Paired-end sequencing (2x 150bp) of four indexed libraries was performed on the Illumina HiSeq2000 platform, generating in excess of 80M mappable reads per sample. Library preparation and sequencing were performed at the Liverpool Centre for Genomic Research (CGR, http://www.liv.ac.uk/genomic-research/). Statistics of the sequencing are shown in **Table S5**.

### Inference of 3’UTR isoforms from 3’-end RNA-seq

Paired-end stranded RNA-seq reads of 150 bp were mapped to the reference rat genome (UCSC, rn5) using TopHat2 (https://ccb.jhu.edu/software/tophat/index.shtml) allowing up to 20 multi-alignments and 2 read mismatches. The extension of the rat 3’ UTR isoform annotation was performed in two steps: 1) by identifying the longest 3’ UTR, and 2) within this longest 3’ UTR, by identifying alternative 3’ UTR isoforms. To find the longest 3’ UTR, nucleotide-level stranded coverage was first obtained for axonal and cell body samples using genomecov from the BEDTools suite (https://bedtools.readthedocs.io/en/latest/). Continuously transcribed regions were next identified using a sliding window across the genome requiring a minimum coverage of 7 reads in more than 80 positions per window of 100 bp; neighbouring regions separated by low mappable regions were merged as described in (30). Expressed fragments were associated with matching strand overlapping 3’UTR using Ensembl Rn5 version 78 (v78). Isolated expressed fragments that did not overlap with any feature were associated with the closest 3’UTR if (1) the 3’UTR was <10kb and (2) there were no intervening annotations. We filtered assigned expressed fragments to exclude potential intragenic transcription, overlapping transcripts, and retained introns as described in (30). If the expressed sequence continued beyond the end of the annotated 3’UTR, we took the sequence as a new 3’ end.

The workflow used to generate input samples for the 3’end-RNASeq data includes two rounds of linear mRNA amplification as described in (29), which leads to accumulation of the reads at the 3’ end of the transcript. Thus, a marked change in the level of coverage in the 3’ to 5’ end direction is expected to occur at the boundaries of alternative 3’ ends within longest annotated 3’ UTR (the read coverage which arises from such experiment looks like the coverage depicted on Figure 1B). To identify alternative 3’ UTR isoforms we smoothed base-level read coverage along longest 3’ UTR using a running median of 150 nt width (corresponds to read length). We then used the R package Segmentor3IsBack (http://cran.r-project.org/web/packages/Segmentor3IsBack/index.html) to identify positions of change-point along the 3’ UTR that are hypothesised to coincide with 3’ ends. The algorithm models the nucleotide read coverage using a negative binomial distribution to first estimate the number of segments via a penalized likelihood criterion (we imposed an upper boundary of 10 segments) and then identifies change-points along the coverage by determining the global maximum of the log-likelihood of a piece-wise constant model. We applied the algorithm to the raw coverage and log2-scaled coverage of both cell body and axon-derived samples. We then merged all 4 annotations (cell body and axon samples, linear and log scale) and clustered 3’ end located within 50 nts distance, selecting the most promoter-distal annotation. We searched the -100 nts to +50 nts region surrounding the 3’ end termini of Ensembl annotated and newly annotated 3’UTR isoforms for 12 canonical and non-canonical PAS motifs (AATACA, ATTAAA, TATAAA, AATATA, AATAGA, AGTAAA, AATGAA, ACTAAA, CATAAA, GATAAA, AAGAAA, and AATAAA) listed in PolyA_database PolyA_DB (http://exon.umdnj.edu/polya_db/v2/) using the matchPattern function from the Biostrings R package (https://www.rdocumentation.org/packages/Biostrings/versions/2.40.2/topics/matchPattern). We tested for the statistical enrichment of the PAS motifs in 3’UTR isoforms using the Fisher’s exact test. A polyadenylation sites atlas was combined from the following sources: 1) poly(A) site annotation (31) build using 3’-end sequencing libraries in human and mouse, lifted from hg19/mm10 to Rn5 using python library CrossMap (http://crossmap.sourceforge.net/); 2) 3’-end sequencing libraries from rat brain and testes (33); 3) 3’end annotation in Ensembl Rn6, RefSeq Rn5 and Rn6, and XenoRefSeq; 4) polyadenylation sites annotations from PolyA_DB and APADB (http://tools.genxpro.net/apadb/). We next compared the percentage of newly annotated 3’ ends recovered from each source and from the compiled polyadenylation site atlas at several intervals from novel 3’ ends.

### 3’UTR isoform quantification and identification of transcripts localized to axons

The number of reads mapped to -500 nts terminal region of each 3’UTR isoform was used to calculate the expression levels. The density of mapped reads in -500 nts terminal region of 3’UTR isoforms is bimodal, with a low-density peak probably corresponding to background transcription, i.e. 3’UTR isoforms of low abundance or 3’UTR isoforms to which reads were spuriously mapped, and a high-density peak corresponding to expressed 3’UTR isoforms. In order to identify 3’UTR isoforms expressed in axons and cell body, a two-component Gaussian mixture was fitted to the data using the R package mclust (https://CRAN.R-project.org/package=mclust). An isoform was called expressed if in both replicates there were less than 5% chance of belonging to the background category or if in at least one replicate there was more than 10% chance of belonging to the expressed category.

### Differential 3’UTR isoforms expression analysis

We focused the analysis on 4,191 tandem pairs of 3’UTR isoforms expressed in the cell body and/or in axonal samples. To identify transcripts displaying a change in the 3’UTR isoform usage between axon and cell body samples, we scored the differences in promoter-proximal to promoter-distal poly(A) site usage:

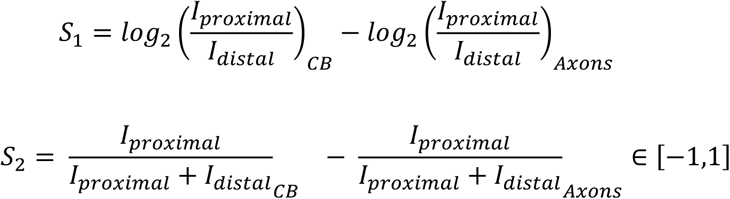

The statistical significance of the changes in proximal-to-distal poly(A) site ratio between cell body and axons was assessed by Fisher’s exact count test using summed-up raw read counts of promoter-proximal versus promoter-distal 3’UTR isoforms originating in the cell body or axonal samples. We applied a False Discovery Rate adjusted threshold of 0.01. A shift towards the usage of promoter-proximal isoforms in axons compared to cell body was considered when S_1 ≤ -1, S_2 ≤ -15% and FDR<0.01. A shift towards the usage of promoter-distal isoforms in axons compared to cell body was considered when S_1 ≥ 1, S_2 ≥ 15% and FDR<0.01. Finally, a stringent threshold was applied to identify highly enriched isoforms in axons as following: for those tandem 3’UTR isoforms showing shift towards the usage of promoter-proximal isoforms in axons as compared to cell body, we required 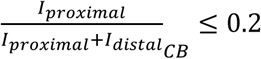. Conversely for those tandem 3’UTR isoforms showing shift towards the usage of promoter-distal isoforms in axons, we required 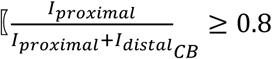.

### Gene Ontology enrichment analysis

GO analysis was performed by comparing pairs of gene lists using the Fisher test with the topGO Bioconductor package (http://bioconductor.org/packages/release/bioc/html/topGO.html). Only GO terms containing at least 10 annotated genes were considered. We applied a *p*-value threshold of 0.05. We manually filtered biologically relevant and statistically enriched GO by removing redundant GO terms and those applying to fewer than 5 genes in the gene lists.

### RT-PCR and quantitative RT-PCR

mRNA was isolated from sympathetic neurons or PC12 cells using TRIzol or RNAEasy mini Kit (QIAGEN) and reverse transcribed with random hexamers and SuperScript III or IV. qRT-PCR reactions (20μL) contained 10μL of Flash SybrGreen Mastermix, or 12.5μL of SybrSelect Mastermix and 0.25μM primers, unless otherwise indicated. Reactions were performed in duplicate or triplicate with the Mastercycler^®^ ep realplex qPCR machine (Eppendorf). For absolute quantification, each experiment included a standard curve, a no-RT control and a no-template control. Standard templates consisted of gel-purified PCR amplicons of known concentrations and each standard curve consisted of seven serial dilutions of the DNA template. For relative quantification, the Comparative Ct Method (ΔΔCt Method) was used. At the end of 40 cycles of amplification, a dissociation curve was performed in which SybrGreen fluorescence was measured at 1°C intervals between the annealing temperature and 100°C. Melting temperatures of amplicons varied between 80°C and 92°C. Primer sequences and PCR conditions are described in Table S4.

### Northern blotting

RNA purified from SCG neurons cultured for 7 days or PC12 cells was separated by electrophoresis in denaturing conditions and transferred to nylon membrane by capillary blotting according to standard protocols. Probes corresponding to IMPA1, Firefly Luciferase or GAPDH coding sequences, or IMPA1L 120nt fragment were labeled using Random Priming Labeling Kits (Takara or Roche) and [α−^32^P] dCTP. Blots were exposed to films or phosphorimager screens and radioactive signal was quantified using ImageJ or ImageQuant TL software (GE Healthcare), respectively.

### Poly(A) tail length assessment

(14). Poly(A) tail length test was performed using USB Poly(A) tail length assay kit following manufacturer’s instructions. Briefly, total RNA was purified from PC12 cells and tagged by G/I tailing to the end of the mRNA using PolyA polymerase (37°C for 60min). The tagged RNA was reverse transcribed (44°C for 60min) using the kit reverse transcriptase and a primer that anneals to the G/I tail. cDNA was then amplified using the IMPA1-1276 or IMPA1-2027 Forward primers, that anneal just upstream of the IMPA1-C or IMPA1-L cleavage site, respectively. Primer sequences and PCR conditions are described in **Table S4**.

### 3’ Rapid Amplification of cDNA Ends (3’ RACE)

Full length 3’UTR of *IMPA1*, *actin beta*, *stathmin 2* and *cofilin1* mRNAs were amplified from axonal and cell body compartments by performing 3’ RACE reactions on total RNA isolated from compartmentalized chambers as previously described (5). Samples were concentrated by speed-vacuum, RNA was divided in two equal samples and used for amplification with SMART RACE cDNA Amplification Kit (Clontech) according to manufacturer’s instructions. Gene specific primers for 3’ RACE assays are listed in **Table S4**. Amplification was performed using Advantage GC 2 PCR kit (Clontech) and PCR products were cloned and sequenced.

### Cloning

*IMPA1 Cleaved* (*IMPA1-C*) and *Long* (*IMPA1-L*) 3’UTR sequences were amplified by PCR from the corresponding RACE clones. After digestion with *NotI*/*XhoI*, IMPA1-C DNA fragment was purified and used to replace IMPA1-L in myrdEGFP-IMPA1-L (5). Mouse IMPA1 coding sequence was PCR-amplified from mouse brain cDNA using primers encoding the HA tag. After digestion with *BamHI*/*NotI, ms* HA-IMPA1 DNA fragments were purified and used to replace dEGFP sequence in myrdEGFP-IMPA1-L or myrdEGFP-IMPA1-C. The 120 nts localization signal of IMPA1-L was cloned by PCR from a IMPA1-L RACE clone and cloned at the 3’ of *ms* HA-IMPA1-IMPA1-C plasmid. To generate Firefly IMPA1-C, -L or –LΔpoly(A) vectors, Firefly luciferase coding sequence was PCR amplified from pGL3 vector (Promega) with primers containing restriction sites for *BamHI* and *NotI*. The DNA fragment was purified and used to replace the myrdEGFP sequence in myrdEGFP-IMPA1-C or myrdEGFP-IMPA1-L. Mutation of IMPA1-C and IMPA-1S poly(A) sites was performed by PCR site-directed mutagenesis (Agilent) of Firefly-IMPA1-L vector. In all Firefly constructs, the bovine Growth Hormone poly(A) sequence was removed by PCR site-directed mutagenesis to create an extra *XhoI* site, that was used for digestion and re-ligation. Primer sequences and PCR conditions are described in **Table S4**.

### Electroporation and analysis of mRNA transport in axons

Neurons were electroporated with the indicated constructs as previously described (http://www.cellectricon.se/pdf/Sympathetic_neurons.pdf). MyrdEGFP was detected by GFP immunostaining. Confocal images were acquired with a SP5 confocal system (Leica) using LAS AF software and automated tiling over several z-stacks, to cover the whole thickness and length of the axons. Maximal intensity projections were processed with Fiji software. Axons were traced manually using NeuronJ plugin and grey value intensity over length was measured. Data analysis was performed using Excel software to calculate average values and standard error means of the intensity for each 200μm axonal segment.

### Quantification of axon degeneration

SGC explants were grown for 36 hours before electroporation with the indicated siRNAs (150nM, GE Dharmacon) and a GFP expression vector (20ng/μL), in the presence of either *ms* HA-IMPA1-C, *ms* HA-IMPA1-C+120, or *ms* HA-IMPA1-L DNAs, (200 ng/μL), as indicated. After 6 days, GFP fluorescence was detected with an inverted Leica epifluorescence microscope, and intact axon bundles that showed no sign of breakdown (i.e. the classical beads-on-string morphology of degenerating axons) were quantified. For imaging, explants were fixed in 4% PFA and stained with anti-GFP and anti-neurofilament antibodies. Antibodies and probing conditions are described in **Table S4**.

### Co-immunoprecipitation and western blotting

Co-immunoprecipitation samples were obtained by lysing cells in RIPA buffer (50mM Tris-HCl pH 7.4, 150mM NaCl, 1% NP-40, 0.5% Sodium deoxycholate, 0.1% SDS, 1mM EDTA, Protease Inhibitors Cocktail) for 10min on ice. After centrifugation, protein concentration in the supernatants was assayed by Pierce^TM^ BCA Assay, and 0.5-1 mg of pre-cleared protein sample was incubated with 2μg of antibody as indicated, overnight at 4°C, on constant rotation. Immuno-complexes were precipitated by adding protein A-agarose beads (GE Healthcare) at 4°C for 2hrs. After extensive washes with RIPA buffer, immunocomplexes were eluted from the beads by boiling in 1X LDS-buffer +2.5% βmercaptoethanol. Samples were resolved on 4-12% PAA pre-cast gels and blotted on PVDF membrane (Amersham). For western blotting, cells were rinsed with PBS and lysed in the plates with 1X LDS-buffer +10% βmercaptoethanol. SDS-PAGE and blotting was then performed as described above. For immunodetection, membranes were blocked in 5% milk for 1hr at room temperature and incubated overnight with the indicated antibodies. Antibodies and probing conditions are shown in **Table S4**.

### RNA ImmunoPrecipitation (RIP)

RNA immunoprecipitation was performed as described (33) with minor modifications. Briefly, protein A/G agarose beads (Santa Cruz) were incubated with antibody (5μg in 1% BSA in PBS) and heparin (1mg/ml) for 2hrs at 4°C, washed with washing buffer (150mM NaCl, 50mM Tris-HCl [pH 8.0], 1% Triton X-100), and incubated with 250-300μg of protein lysates 1 hr at 4°C. Beads were extensively washed, and RNA was eluted in 0.2M Na Acetate, 1mM EDTA, and 0.2% SDS for 5 min at 70°C. For normalization, 20pg of in vitro transcribed RNA synthesized from the T7 control DNA Template (AmpliScribe™ T7 Transcription Kit, Epicentre) was added to the samples. RNA from inputs and immunocomplexes was purified, subjected to DNAse digestion (Ambion), reverse transcribed and assayed by qPCR. Primer sequences and PCR conditions are described in **Table S4**.

### Dual luciferase assay

PC12 cells were transfected with the indicated Firefly Luciferase-IMPA1 constructs and thymidine-kinase – Renilla Luciferase (Promega) using Lipofectamine 2000 for 48hrs. Samples were processed using the dual-luciferase reporter assay system (Promega), according to manufacturer’s instructions.

### Polysome fractionation

Polysome fractionation was performed as described (34). Briefly, PC12 cells were lysed in ice-cold gradient buffer (0.3M NaCl, 1mM MgCl2, 15mM Tris-HCl (PH7.4), 0.1mg/mL cyclohexamide and 1mg/mL heparin, 1% Triton X-100, 500U/mL RNAse inhibitors). Samples were centrifuged and the supernatants layered onto 10–50% sucrose linear gradients. The gradients were sedimented at 38,000 r.p.m., using a SW40Ti rotor (Beckman) or a Sorvall TH-641 rotor for 2 hrs at 4°C. Eleven fractions (1mL each) were collected from the gradients and transferred in 3ml of 7.7M guanidine-HCL using a Foxy R1 gradient fractionator (Teledyne ISCO; ISCO peak Trak version 1.10 software) with continuous measurement of the absorbance at 254nm. RNA was precipitated, treated with DNAse and purified using RNAeasy Mini Kit (QIAGEN). For fractions 1 and 2, protocol was modified as suggested by manufacturer for recovery of small size RNA. Samples were concentrated by speed-vacuum and analysed by northern blot.

### Mass spectrometry

Immuno-complexes were precipitated from 20x10^6^ PC12 cells naïve or differentiated with NGF for 4 days. On-bead digestion and nano LC-MS/MS analysis was performed as described (35) with minor changes. The procedure is summarized in **Fig. S6** and described briefly below. Immuno-precipitated proteins were released from the resin by on-beads digestion for 15min at 37°C using 200ng of trypsin (Promega). The supernatants were collected and subjected to conventional in-solution tryptic digestion (overnight at 37°C) in denaturing conditions (reduction by 10mM DTT for 1hr at 37°C followed by 24mM iodoacetamide for 1hr at 37°C quenched by addition of 2mM DTT for 30min at 37°C). Tryptic peptides were then subjected to differential labelling by either oxygen^18^ or dimethyl labelling. Pairs of differently labelled samples were mixed, purified by StageTips and subjected to nano LC-MS/MS analysis. Chromatography was performed on an Easy LC 1000 nanoLC system (Thermo Fisher Scientific, Odense, Denmark). The analytical nanoLC column was a pulled fused silica capillary, 75μm i.d., in-house packed to a length of 10 cm with 3μm C18 silica particles from Dr. Maisch GmbH (Entringen, Germany). A 60-min binary gradient was used for peptide elution. MS detection was performed on a quadruple-orbitrap mass spectrometer Q-Exactive (Thermo Fisher Scientific) operating in positive ion mode and data-dependent (Top-12) scanning mode. Data were processed using Proteome Discoverer 1.4 (Thermo Fisher Scientific), using Sequest as search engine, and querying the March 2015 RATTUS reference proteome sequence database (http://www.ebi.ac.uk/uniprot). The protein sequence database was merged with a list of common contaminants named “Common Repository of Adventitious Proteins” retrieved from The Global Proteome Machine website (http://www.thegpm.org/crap/index.html). In total, 27,927 entries were searched. Peptide identifications were validated by Percolator integrated in Proteome Discoverer (http://www.matrixscience.com/help/percolator_help.html). Percolator q-value was set to equal or less than 0.05. Quantification values based on < 3 peptides were manually checked in raw MS data. MS/MS data relative to protein hits identified by a single peptide are reported in **Table S3**. Protein H:L ratios obtained from all technical replicates of a given biological replicate were transformed into log2 space before their median was calculated.

### RNA oligo-Mediated Ligation (RML) RT-PCR

RML RT-PCR was performed as described (36) with the following modifications. Cleaved fragments were isolated and cloned using 1.2ng of axonal RNA purified and pooled from 55 explants where the cell bodies had been surgically removed, or 1.5 μg or less of total cellular RNA. Total cellular RNA was DNAse-digested and purified by phenol:chloroform purification. Quality control of starting material was performed using Agilent Tapestation 2200 (UCL Genomics). Samples with a RIN value≥ 7.2 were used for RLM RT-PCR. RNA was denatured and tagged by ligation with 25ng or 250ng for axonal or total RNA, of RNA oligo and 30U of T4 RNA ligase (NEB) for 1hr at 37°C followed by overnight incubation at 16°C in a PCR machine. Ligated axonal RNA was then purified using buffer PB (QIAGEN) +10% β−mercaptoethanol and AMPure XP beads (Beckman Coulter) as per manufacturer’s instructions. Ligated RNA was then reverse transcribed using random hexamers and 50U SuperScript IV reverse transcriptase for 1hr at 50°C. After RNAseH (NEB) digestion, cleaved fragments were amplified by PCR using Q5 DNA polymerase (NEB) and cloned in pCR™4Blunt-TOPO^®^ vector according to manufacturer’s instruction. At least 17 individual, random clones were analyzed by sequencing. When used for RT-qPCR, amplification of the cleaved fragments was carried out in 25μL reaction using SybrSelect MasterMix. Primer sequences and PCR conditions are described in **Table S4**.

### Radioactive *in vitro* cleavage assay

Assay were performed as described with the following modifications (11, 36). RNA oligos were prepared by *in vitro* transcription using mirVana probe construction kit according to manufacturer’s instruction. RNA was then dephosphorylated using Calf Intestine Phosphatase (NEB) and purified by phenol:chloroform extraction. After precipitation, RNA probes were labelled at the 5’ using [γ-^32^P] ATP and T4 polynucleotide kinase. After gel purification of full size probes, oligos were incubated for 2 hrs at 26°C with cytoplasmic protein fractions prepared from sympathetic neurons using NE-PER kit (Pierce). *In vitro* cleavage assays were performed by adding 50nM human recombinant Ago2 (expressed in baculovirus, Active Motif) in a reaction mixture containing 25mM Hepes-KOH pH 7.5, 50mM KOAc, 5mM Mg(OAc)_2_, 5mM DTT for 1.5 hrs at 26°C. When testing Ago2 biological activity, 30nM recombinant Ago2 was incubated for 2 hrs at 26°C with 30nM single-stranded, phosphorylated *luc* guide siRNA prior to cleavage assay of *Luc* miRNA-target oligo. Following purification using Triazol, samples were separated on 8% acrylamide gel in denaturing conditions and gels were exposed to X-rays.

### Statistical analyses

Data are expressed as averages ± SEM. One-way ANOVA with post-hoc test or t-test were used as indicated to test for statistical significance, which was placed at *p* < 0.05 unless otherwise noted. In all experiments, *n* refers to independent biological replicates from independent cell cultures.

### Accession numbers

The RNA-seq data and the new 3’UTR annotation will be available at Gene Expression Omnibus; the accession number is in preparation.

## Acknowledgments

We thank Cristina Ottone for generating the Firefly-IMPA1 3’UTRs constructs, and Carola Zimmermann for sharing the drawing and the staining of compartmentalized chambers, and for performing the Stahmin2 RACE shown in **Fig.S3B**. We also thank Aniko Ludanyi for sharing the results of her screening of RNA-binding protein regulated by NGF. We are indebted to Anne Willis and the Genomic Service (University of Leicester) for the use of the Foxy R1 gradient fractionator and to Tina Daviter (ISMB Biophysics Centre at Birkbeck, University of London) for the use of the phosphorimager. We thank Adolfo Saiardi, Paolo Salomoni and Jernej Ule for insightful suggestions on the manuscript, and all members of the Riccio lab for helpful discussions.

## Data and materials availability

All materials used in the analysis are available upon request. The RNA-seq data and the new 3’UTR annotation will be available at Gene Expression Omnibus; the accession number is in preparation.

---

**Fig. S1.**
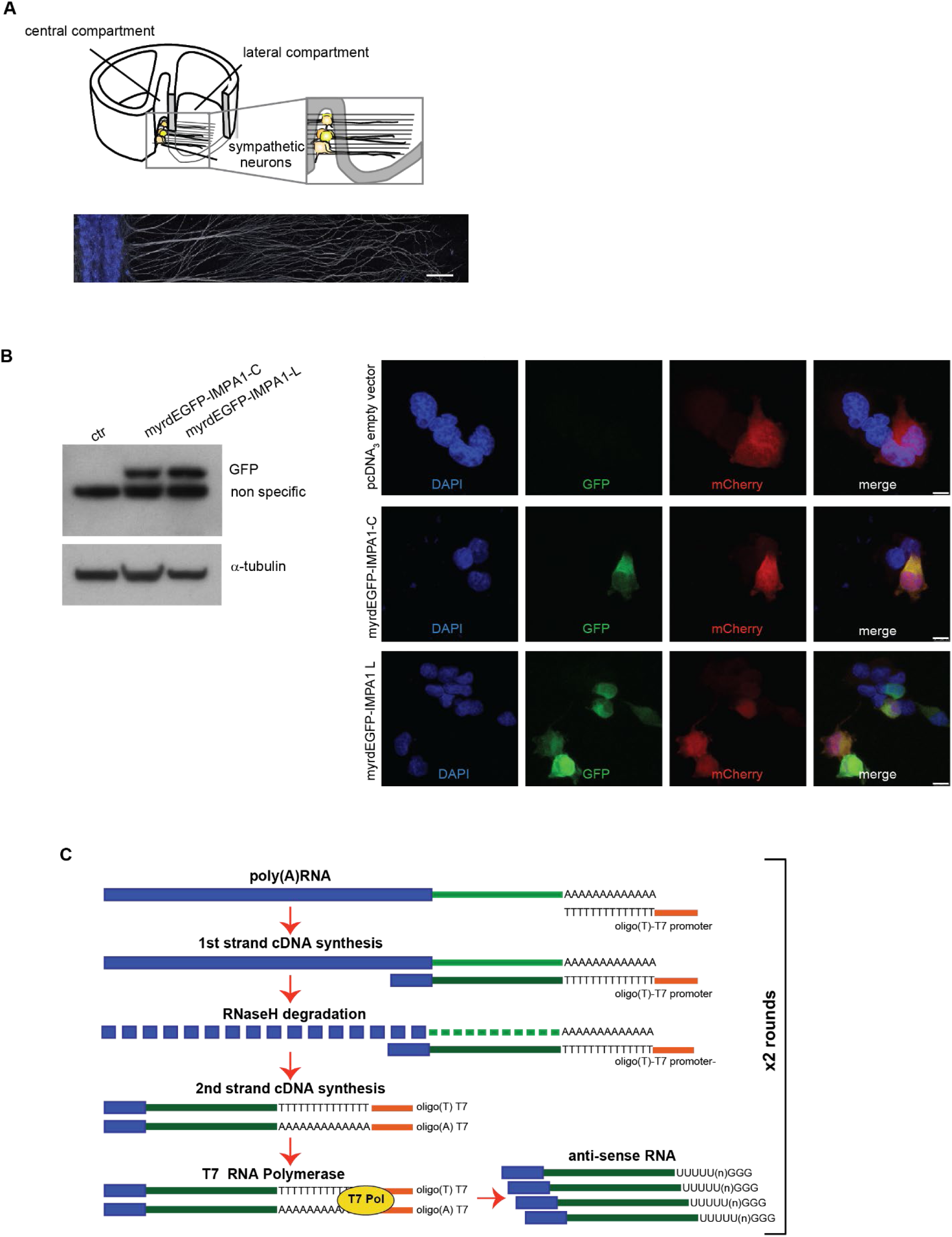
Identification of unique 3’UTR isoforms in axons of sympathetic neurons. (**A**)(*Upper panel*) Schematic representation of a compartmentalized chamber. Neurons are plated in the central compartment and axons grow into the lateral compartments. (*Lower panel*) Immunofluorescence of sympathetic neurons grown in compartmentalized chambers for 10 days and stained with DAPI (blue) and β-tubulin (white) antibody. Scale bar=500μm. (**B**) (*Left*) Western blot analysis of GFP and α-tubulin on PC12 cells transfected with either myrdEGFP-IMPA1-C or myrdEGFP-IMPA1-L. Ctr: non transfected cells. (*Right)* Representative images of naïve PC12 cells co-transfected with empty vector pcDNA3 or myrdEGFP-IMPA1-C or myrdEGFP-IMPA1-L, and mCherry plasmids. The non-specific band detected by western blotting does not affect the immunofluorescence staining. Scale bar 75μm. (n=3). (**C**) Workflow of linear amplification of mRNA. (See also **Fig. 1**).

**Fig. S2.**
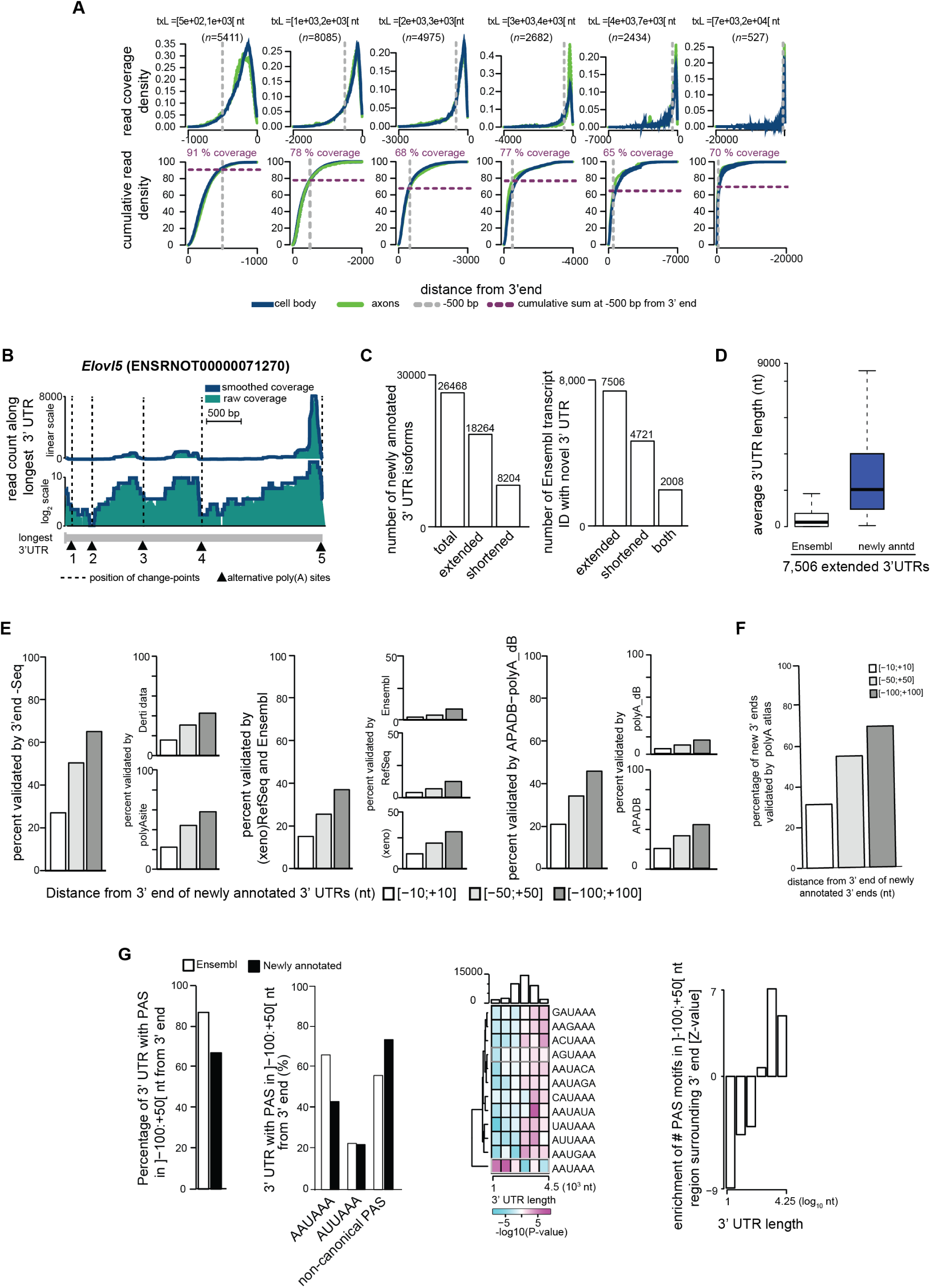
Analysis of 3’UTR PAS choice in axons and cell bodies of sympathetic neurons. (**A**) Reads accumulation at 3’ end of the Ensembl transcripts in function of transcript length. (*Upper*) Read density coverage and (*Lower*) cumulative read density along transcript are shown. (**B**) Example of identification of novel 3’ ends in the longest 3’UTR of Elovl5. Raw coverage was smoothed using a running median (window width = 100 nts) and potential 3’ ends were identified by segmenting sudden transitions in read depth. (**C**) (*Left*) Number of newly annotated 3’UTR isoforms compared with Ensembl Rn5 annotations. (*Right*) Number of Ensembl transcript ID expanded with newly annotated 3’UTRs. (**D**) Average length of the 3’UTRs of 7,506 Ensembl transcript ID extended by intersecting expressed genomic fragments with Ensembl Rn5 annotation. (**E**) Percentage of 3’UTR isoforms for which the indicated region surrounding the 3’ end intersects with a PAS obtained from 3’-Seq data, a 3’ termini annotated in RefSeq (Rn5, Rn6 and XenoRefSeq) or Ensembl (Rn6), or a PAS annotated in APADB or PolyA_DB2. Comparison between combined (*Left*) and individual (*Right*) datasets is shown in each panel. (**F**) Percentage of newly annotated 3’ ends recovered from a polyadenylation site atlas compiled from multiple sources (see Experimental Procedures) at the indicated distance intervals from novel 3’ ends. (**G**) (*Left*) Frequency of canonical and variant PAS motifs detected between -100 to +50 nt of newly annotated (black) or Ensembl annotated (white) 3’ ends. Total PAS motifs and canonical vs. variant PAS motifs are shown. (*Middle*) Relative occurrence of different PAS motifs in promoter-proximal and promoter-distal 3’UTRs. Upper column graph indicates the number of 3’UTR isoforms per range of 3’UTR length. Color-scale: -log10(P-value) of enrichment in PAS motif obtained by Fisher test of the number of 3’ end that contains at least one motif per range of 3’UTR length. (*Right*) Relative occurrence of PAS motifs in the [-100;+50] nt region surrounding the 3’ ends at increasing length. (See also **Fig. 1**).

**Fig. S3.**
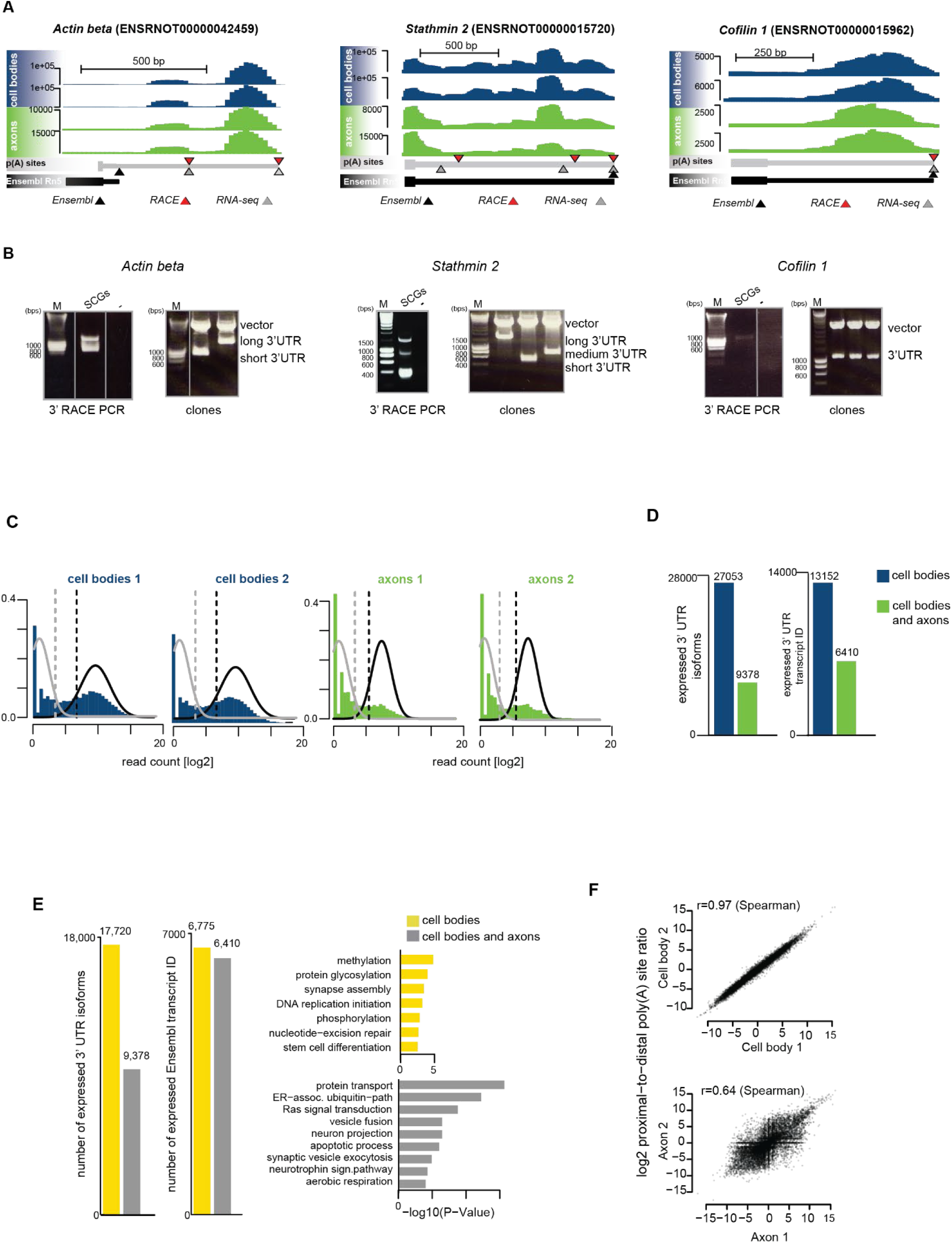
Analysis of 3’UTR PAS choice in axons and cell bodies of sympathetic neurons. (**A**) Genome browser view of the Actin beta, Stathmin 2 and Cofilin 1 3’UTRs. 3’ end isoforms annotated in Ensembl Rn5 or identified by RNA-seq data and by RACE are indicated by arrowheads. (**B**) (*Left panels*) Agarose gel analysis of RACE products to amplify *actin beta*, *Stathmin 2* and *Cofilin 1* 3’UTRs and (*Right panels*) of restriction digestions of representative clones obtained by cloning of corresponding RACE PCR products. (**C**) Identification of 3’UTR isoforms expressed in cell bodies (blue) and axons (green) performed by fitting bimodal distribution on log2-raw count mapping the 500 nts distal region of 3’ end. (**D**) Number of 3’UTR isoforms (*Left*) and Ensembl transcript ID (*Right*) expressed in cell bodies and axons. (**E**) (*Left*) Comparative analysis of 3’UTR isoforms and transcript IDs enriched in cell bodies only (yellow) or cell bodies and axons (grey). (*Right*) Statistically enriched GO terms of genes identified in cell bodies only or cell bodies and axons samples. (**F**) Scatter plots of the relative use of promoter-proximal and promoter-distal poly(A) sites in two biological replicates of cell body (Upper) and axon (Lower) samples. (See also **Fig. 1**).

**Fig. S4.**
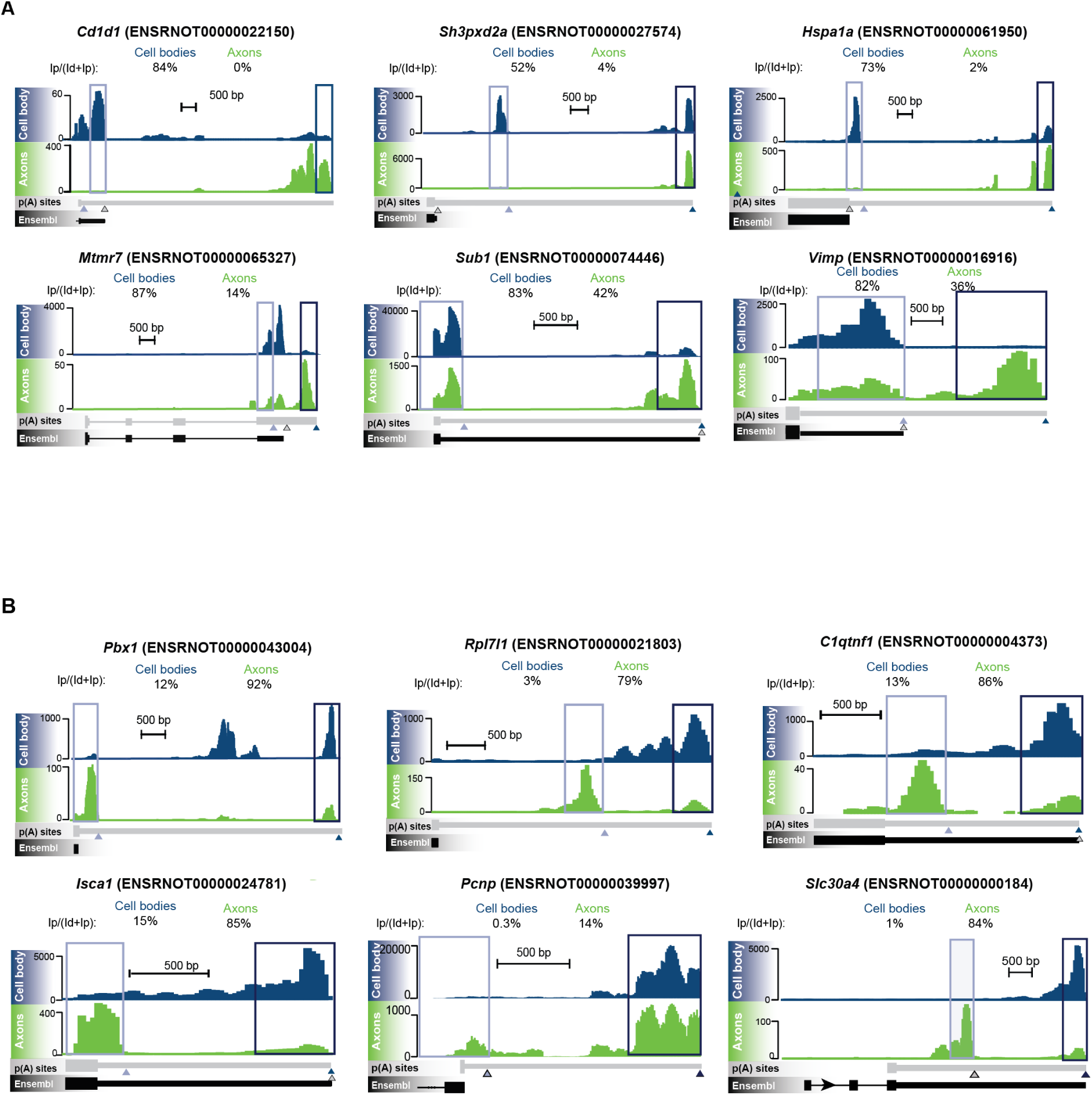
Axonal transcripts with a proximal or distal 3’UTR bias identified by 3’end RNAseq. (A and B) Examples of transcripts with a marked shift towards (**A**) increased promoter-distal poly(A) site usage or (**B**) increased promoter-proximal poly(A) site usage, in axons compared to cell bodies. (See also **Fig. 1**).

**Fig. S5.**
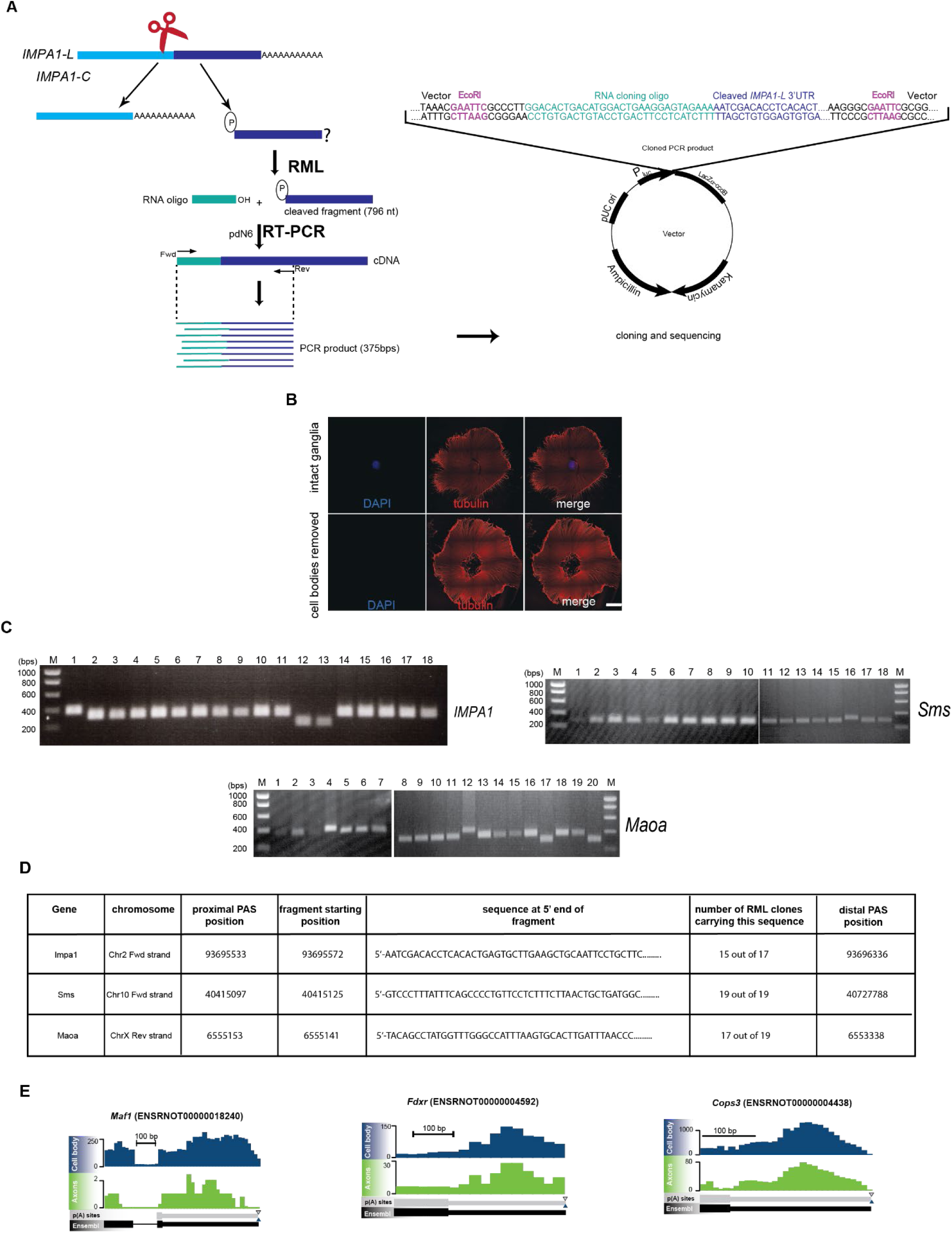
Analysis of axonal cleavage of transcripts by RML-RT-PCR. (**A**) Schematic representation of 5’P-dependent RNA oligo-Mediated Ligation (RML) and cloning experiments. (**B**) α-Tubulin immunostaining of representative SCG explants before (*Top*) or after (*Bottom*) the removal of cell bodies. Nuclei were stained with DAPI. Scale bar = 200 μm. (**C**) *EcoRI* restriction digestions of clones carrying an insert corresponding to IMPA1-L (*Left*), Sms (*Right*) and Maoa (*Bottom*) cleaved fragments in axons. *EcoRI* sites flank the PCR product insertion site for excision of the insert. (**D**) Genomic coordinates (Ensembl Rnor_6.0) for the proximal and distal PAS, and for the 5’ends of the cleaved fragments of IMPA1, Sms and Maoa are listed, together with the sequence at the 5’ end of the cleaved fragments as obtained by sequencing of the indicated number of clones. (**E**) Genome browser view of Maf1, Fdxr and Cops3 transcripts that express single 3’UTR isoforms. (See also **Fig. 2A, B**).

**Fig. S6.**
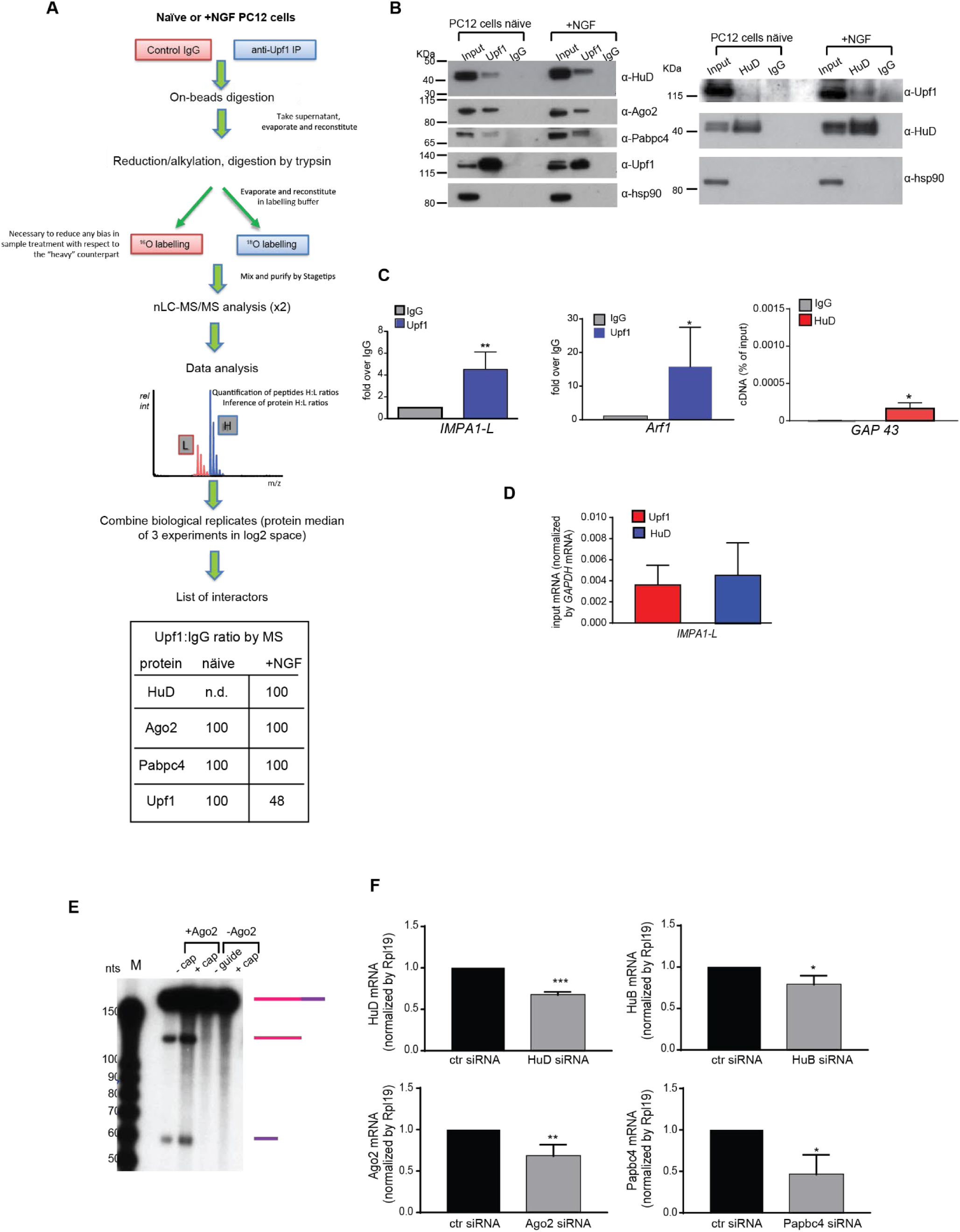
Remodelling of IMPA1 3’UTR depends on a complex that includes Ago2, HuD and Upf1. (**A**) Workflow employed for the discovery of Upf1 interactors in naïve and NGF-stimulated cells. The table shows the enrichment of the indicated interactors in the Upf1 immuno-precipitates of naïve or differentiated (+NGF) PC12 cells as measured by mass-spectrometry. (**B**) Western blotting of Upf1 (*Left*) or HuD (*Right*) co-immuno-precipitated proteins in extracts of naïve and NGF-differentiated PC12 cells. (**C**) (*Left*) Binding of IMPA1-L mRNA to Upf1 as assessed by RIP and quantified by RT-qPCR. Control RIP of Arf1 (*Middle*) or Gap43 (*Right*) mRNAs immunoprecipitated with normal IgG and Upf1 or HuD antibody, respectively, and quantified by qRT-PCR in sympathetic neurons lysates. Data presented as averages ±s.e.m. (*p<0.05; unpaired one-tail t-test n=4-5). (**D**) Normalized expression levels of IMPA1-L mRNA in HuD or Upf1 RIP inputs quantified by RT-qPCR. Data presented as averages ±s.e.m. (**E**) *In vitro* cleavage of luc target RNA by recombinant Ago2 and luc guide siRNA, showing biological activity of the recombinant Ago2 protein. Two fragments of the expected sizes (125 and 57 nts) are detected only in the samples containing Ago2 and guide siRNA. The lack of other fragments demonstrates that the preparation of recombinant Ago2 is devoid of contaminant RNAses. (**F**) RT-qPCR to evaluate efficiency of HuB, HuD, Ago2 and Pabpc4 mRNA silencing. Data presented as averages ±s.e.m. (*p<0.05, **p<0.005, ***p<0.0005; paired t-test, n=4-6). (See also **Fig. 2E,F** and **Fig. 3**).

**Fig. S7.**
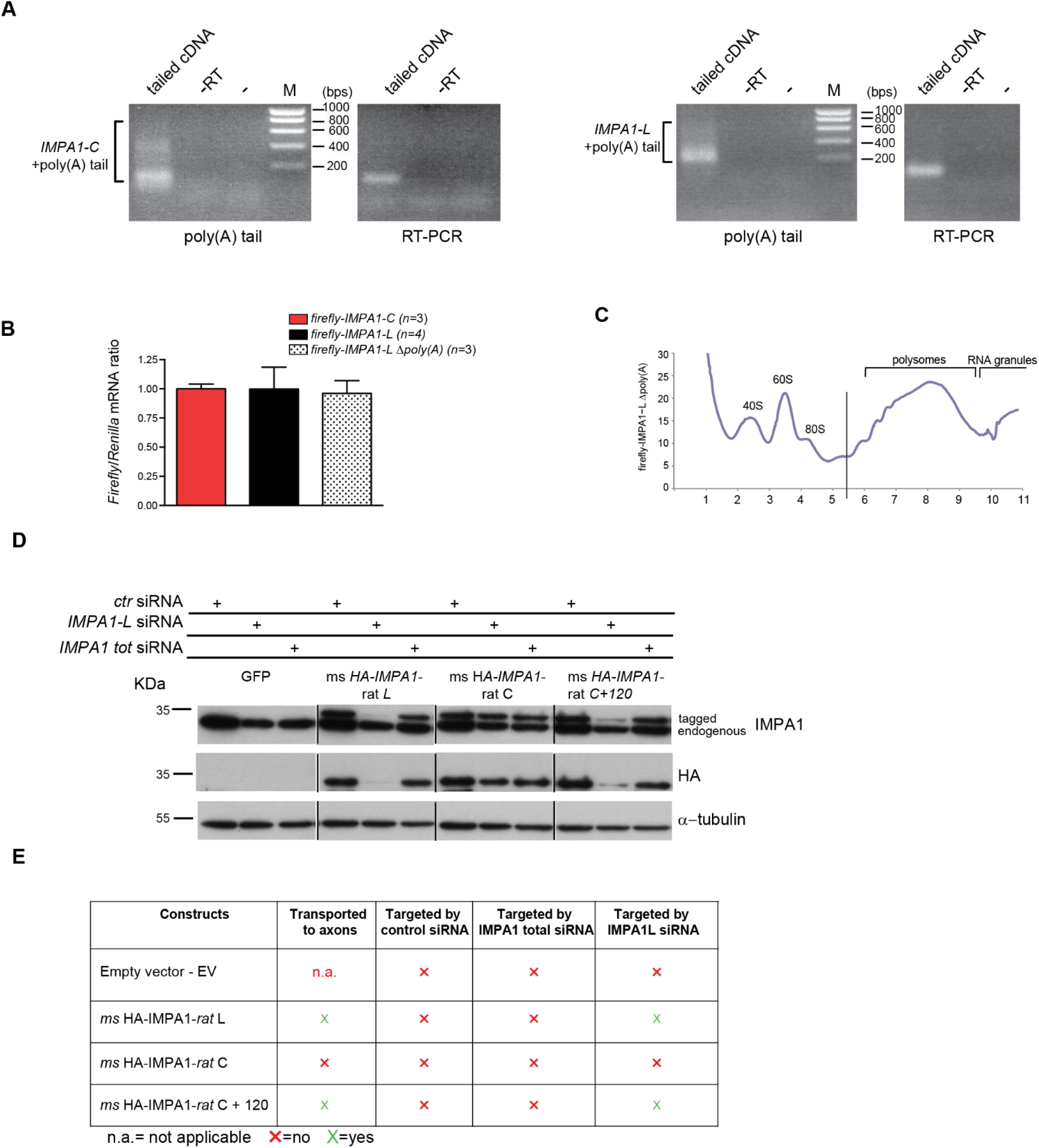
IMPA1-L is transported to axons and cleaved to increase translational efficiency. (**A**) mRNA isolated from PC12 cells was subjected to G/I tailing at the 3’end before assaying poly(A) tail length of IMPA1-C or IMPA1-L 3’UTR by RT-PCR. (Representative experiment of *n*=3). (**B**) qRT-PCR analysis of Firefly and Renilla luciferase mRNAs isolated from PC12 cells transfected with the indicated vectors. Data presented as average ±s.e.m. (**C**) Representative absorbance profile (A254nm) of polysomal fractions isolated from PC12 cells transfected with Firefly-IMPA1L-Δpoly(A). Peaks representing the 40S, 60S and 80S ribosomal subunits, polysomal fractions and RNA granules are indicated. Line shows separation between free-monosomal and polysomal fractions. (**D**) IMPA1, hemagglutinin (HA) and α-tubulin western blotting of PC12 cells transfected with the indicated siRNAs and vectors encoding HA-tagged mouse IMPA1 flanked by either the Long, the Cleaved or the Cleaved + IMPA1-L localization signal 3′ UTRs of rat IMPA1 (ms HA-IMPA1-IMPA1L, ms-HA IMPA1-IMPA1-C and ms HA-Impa1-IMPA1-C+120, respectively). Higher band in top panels is HA-tagged mouse IMPA1 and lower band is endogenous rat IMPA1 (*n*= 3). (**E**) Table summarizing the subcellular localization and silencing of the vectors used in **S7D** and in Fig. 4. (See also **Fig. 4**).

**Fig. S8.**
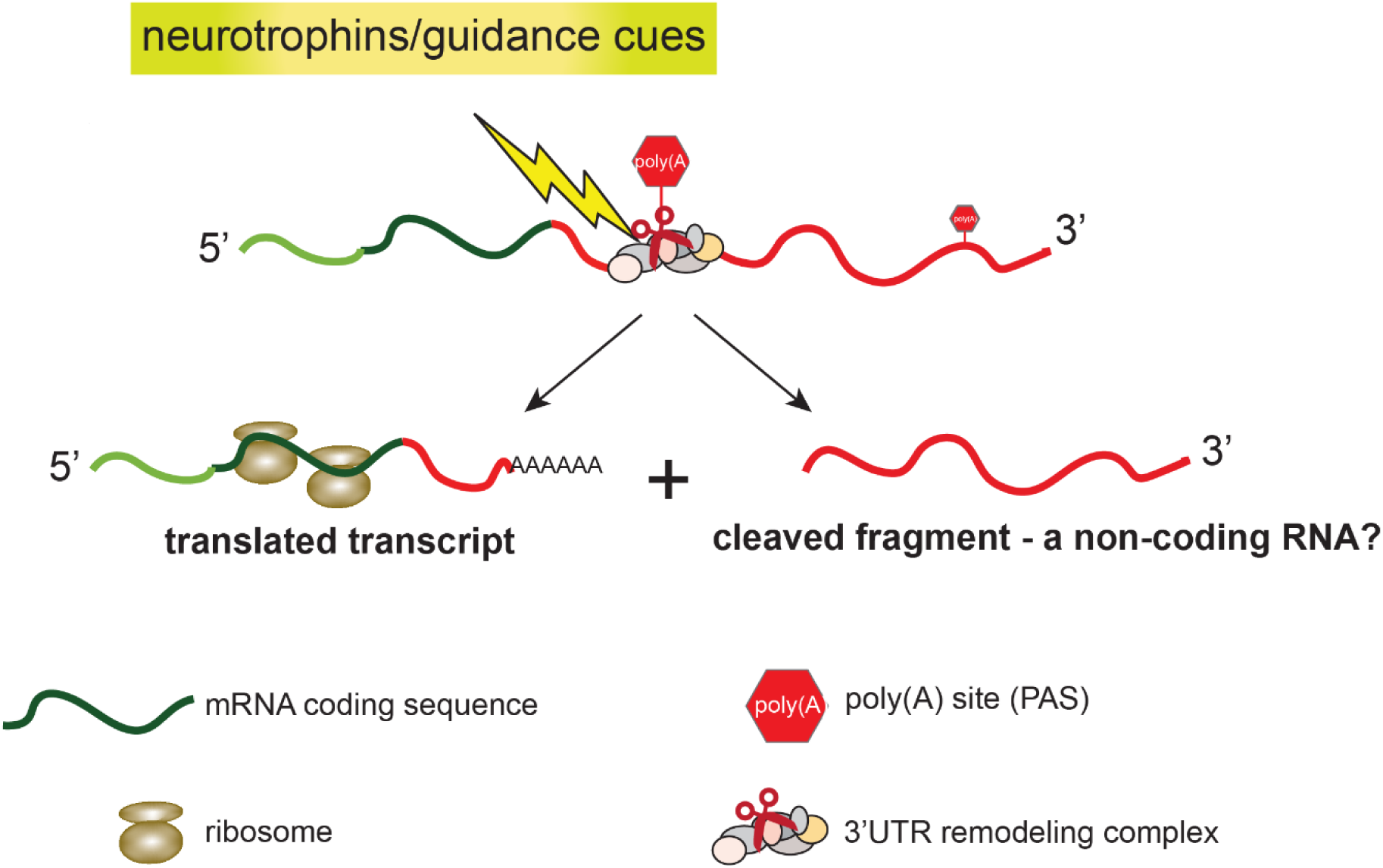
Model depicting the cleavage of transcripts in axons. Schematic representation summarizing the cleavage and remodelling of 3’UTR in axons.

**Table S1.**
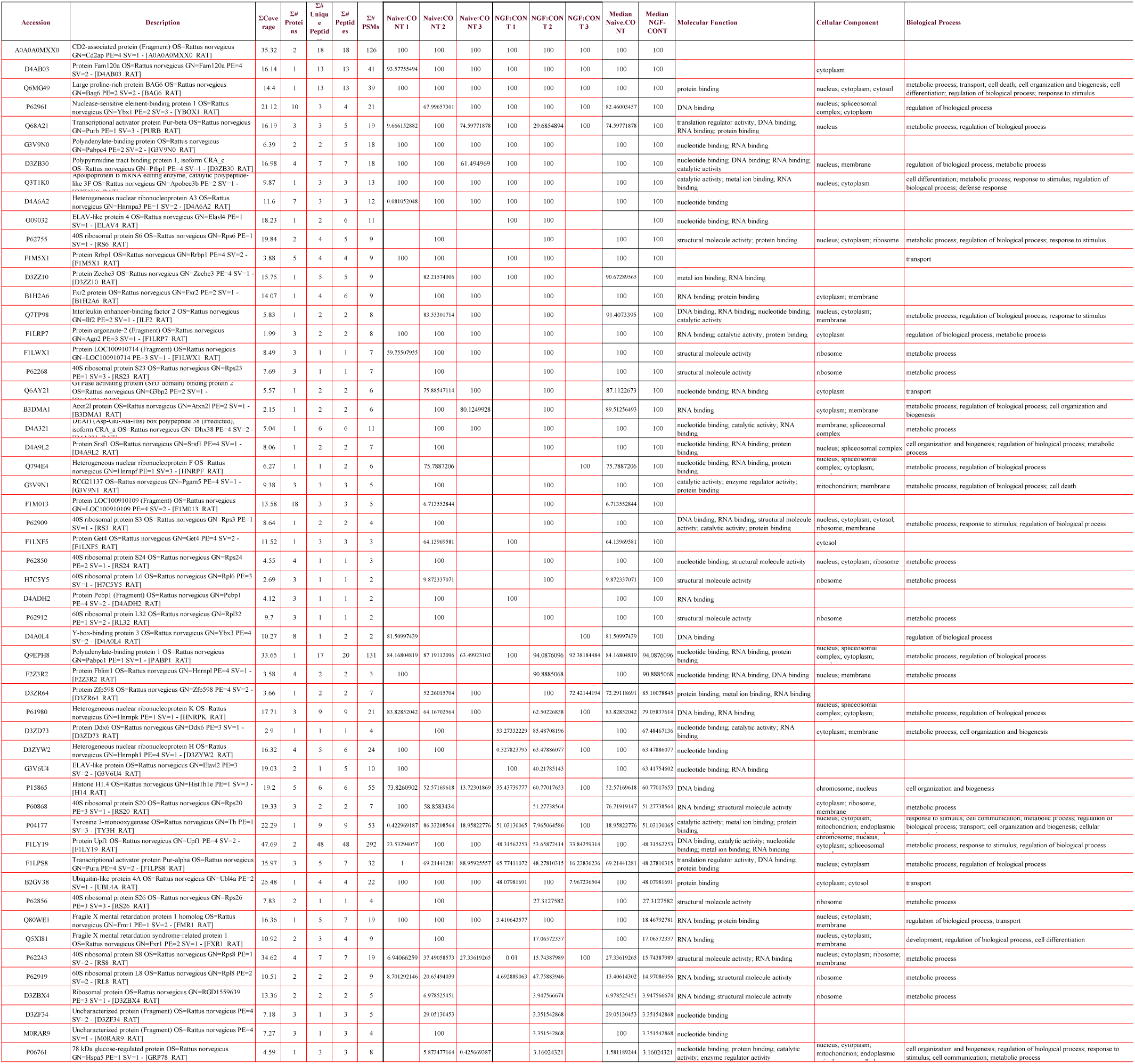
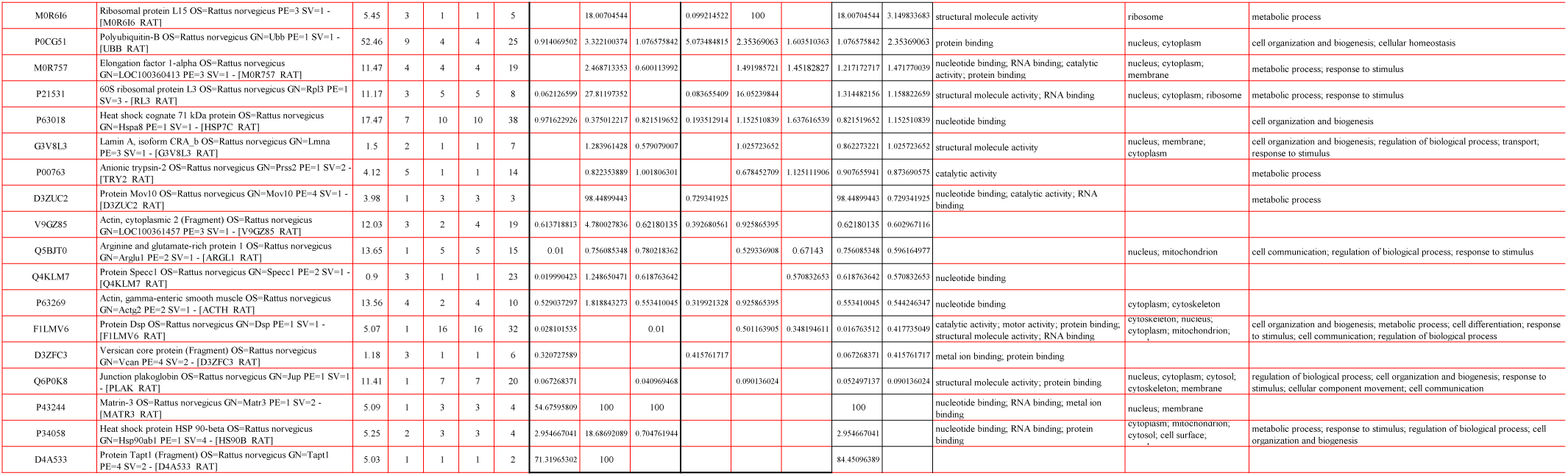
Related to Fig. 2 and S6. List of proteins identified by mass-spectrometry in immunoprecipitation experiments.

**Table S2.**
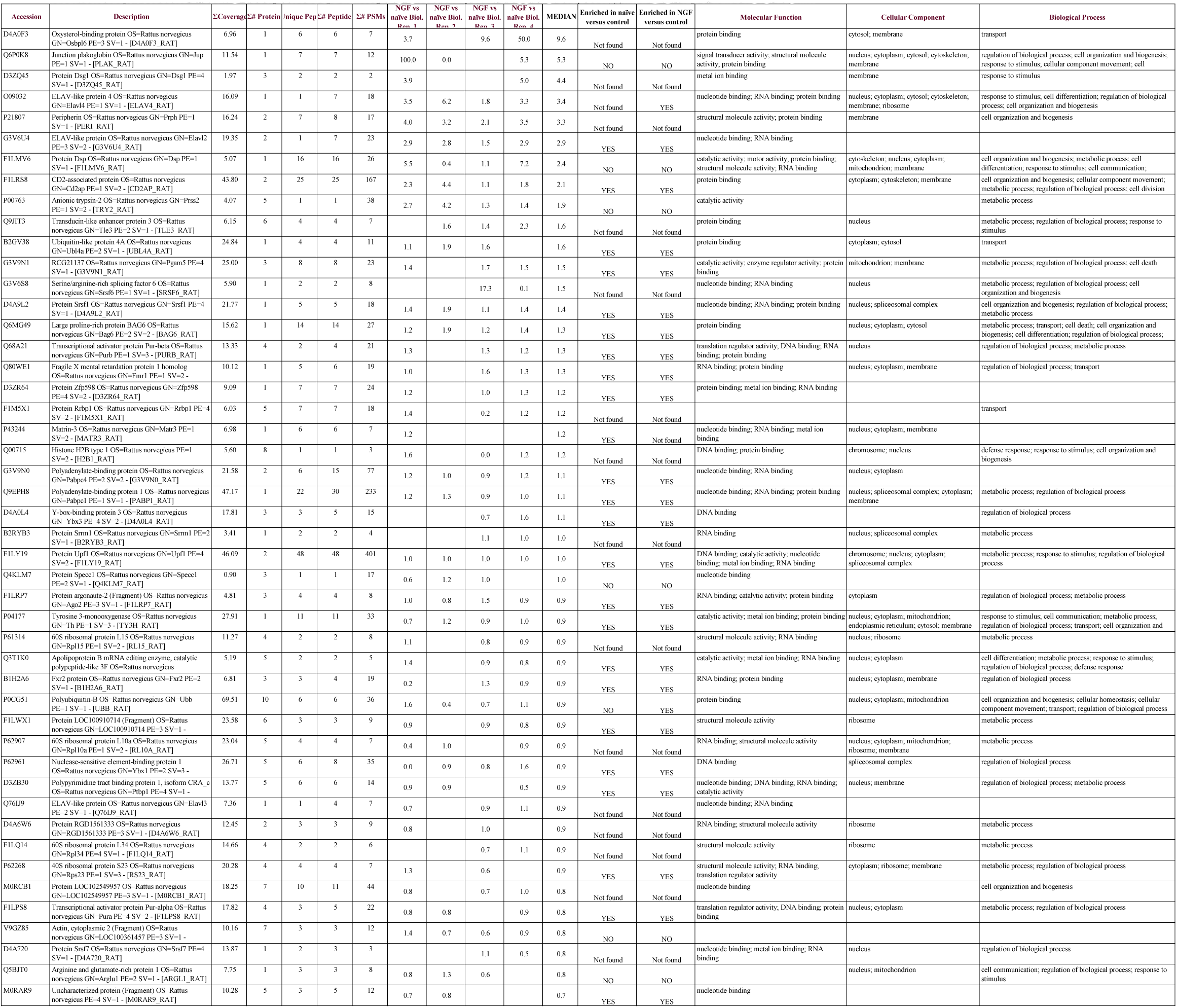
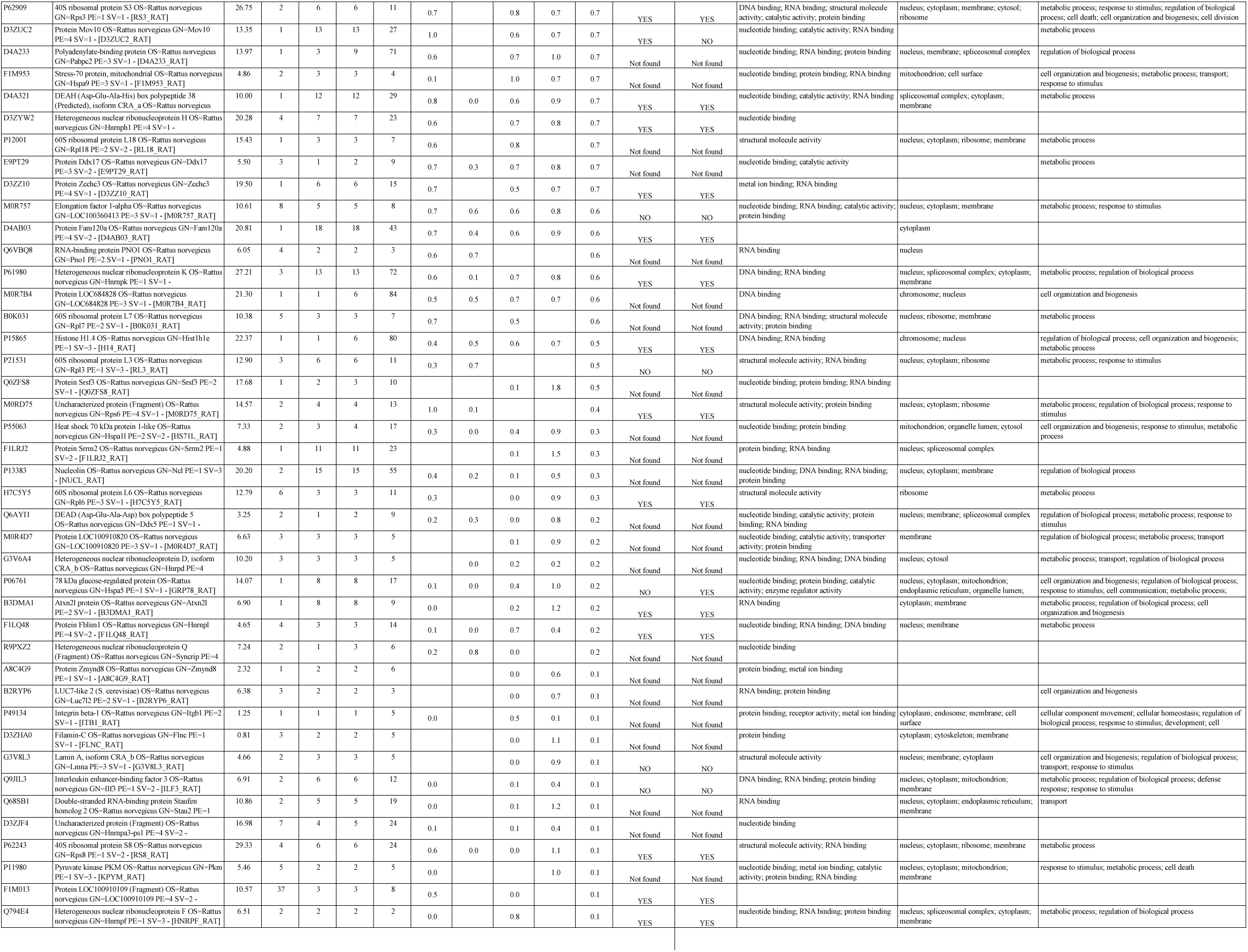
Related to Fig 2 and S6. Putative interactors of Upf1 were singled out according to the following criteria: (i) identified and quantified in at least two replicate experiments; (ii) fold change Upf1 IP:IgG control >2 in all available replicates

**Table S3.**
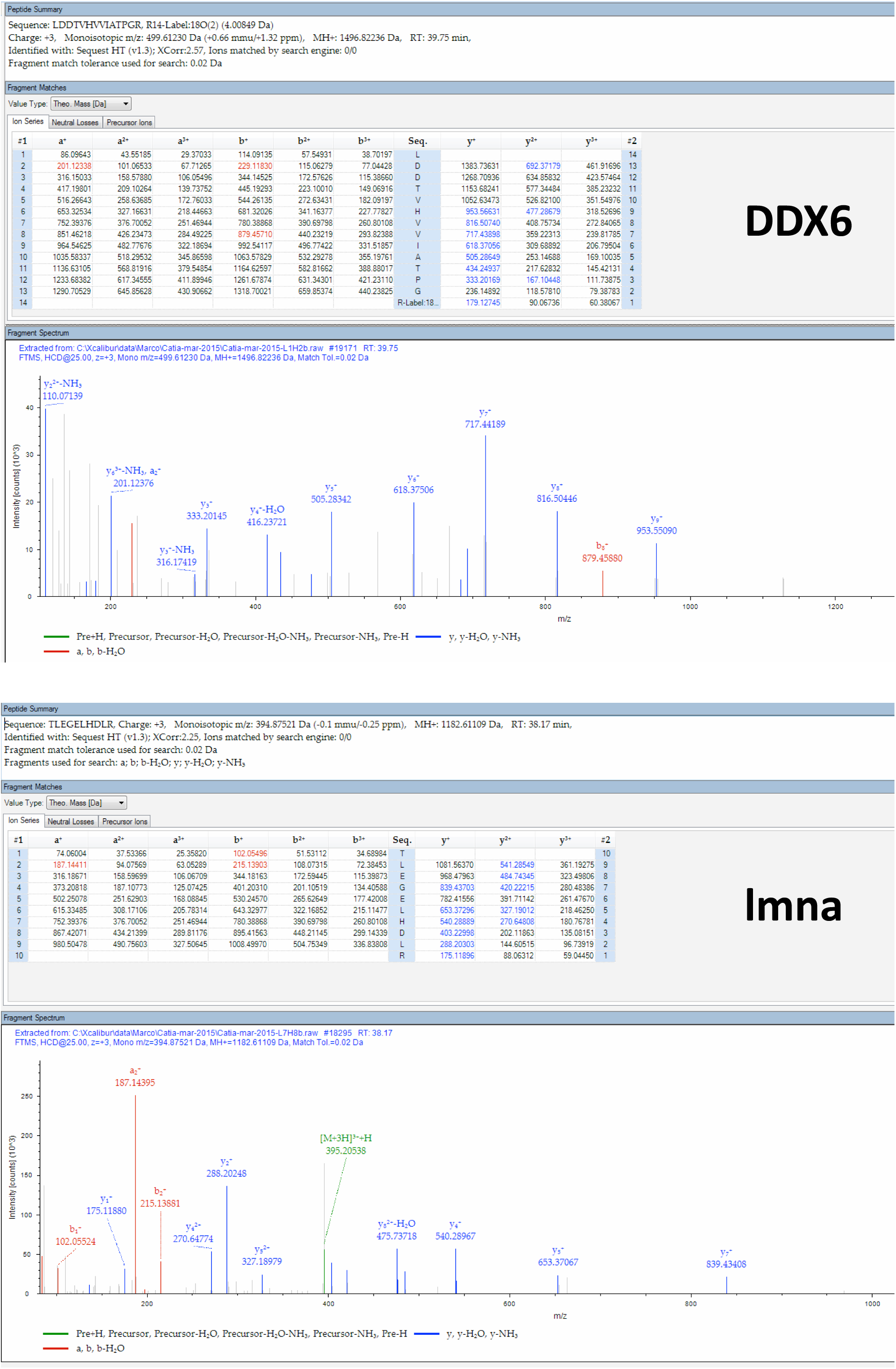
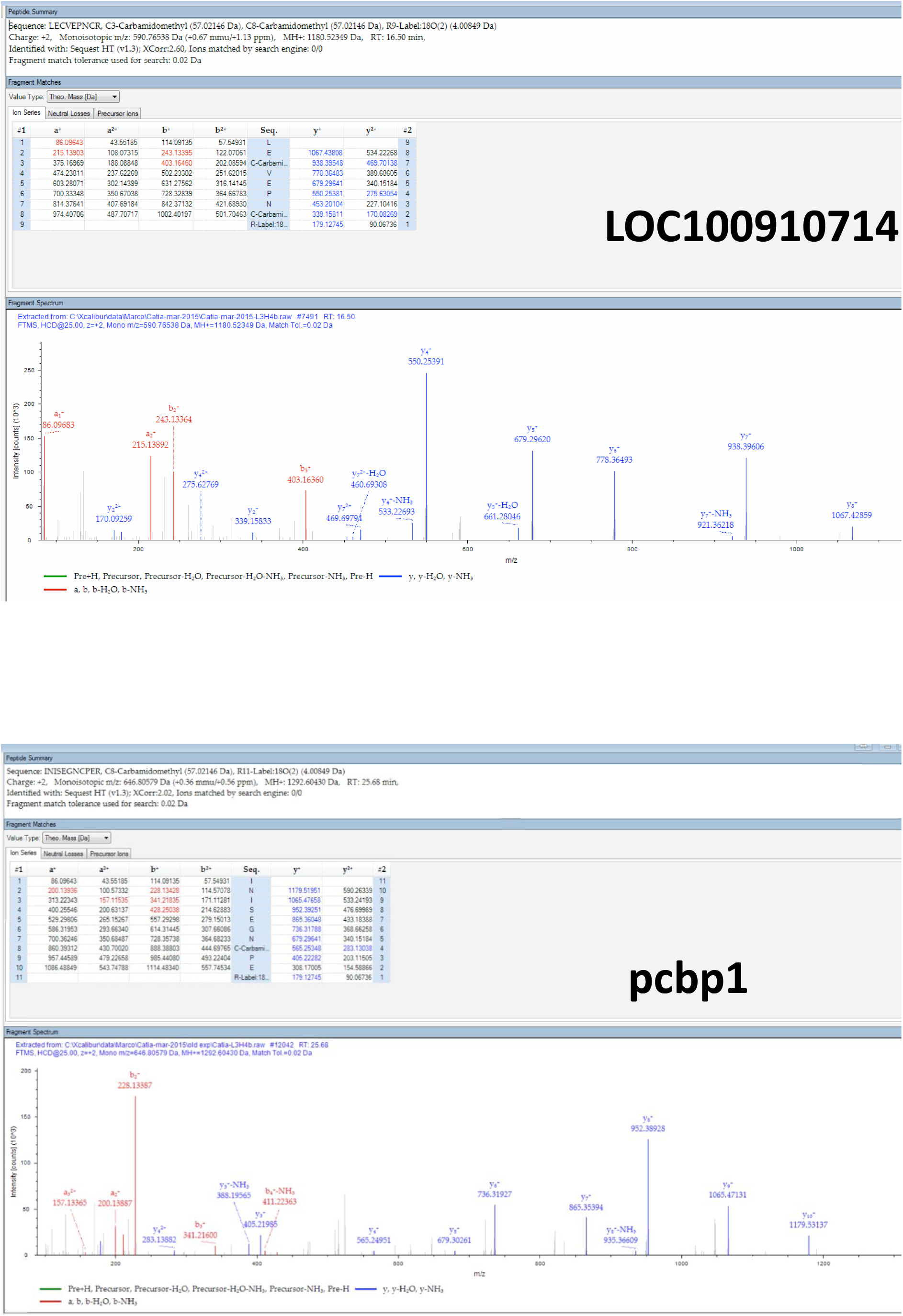
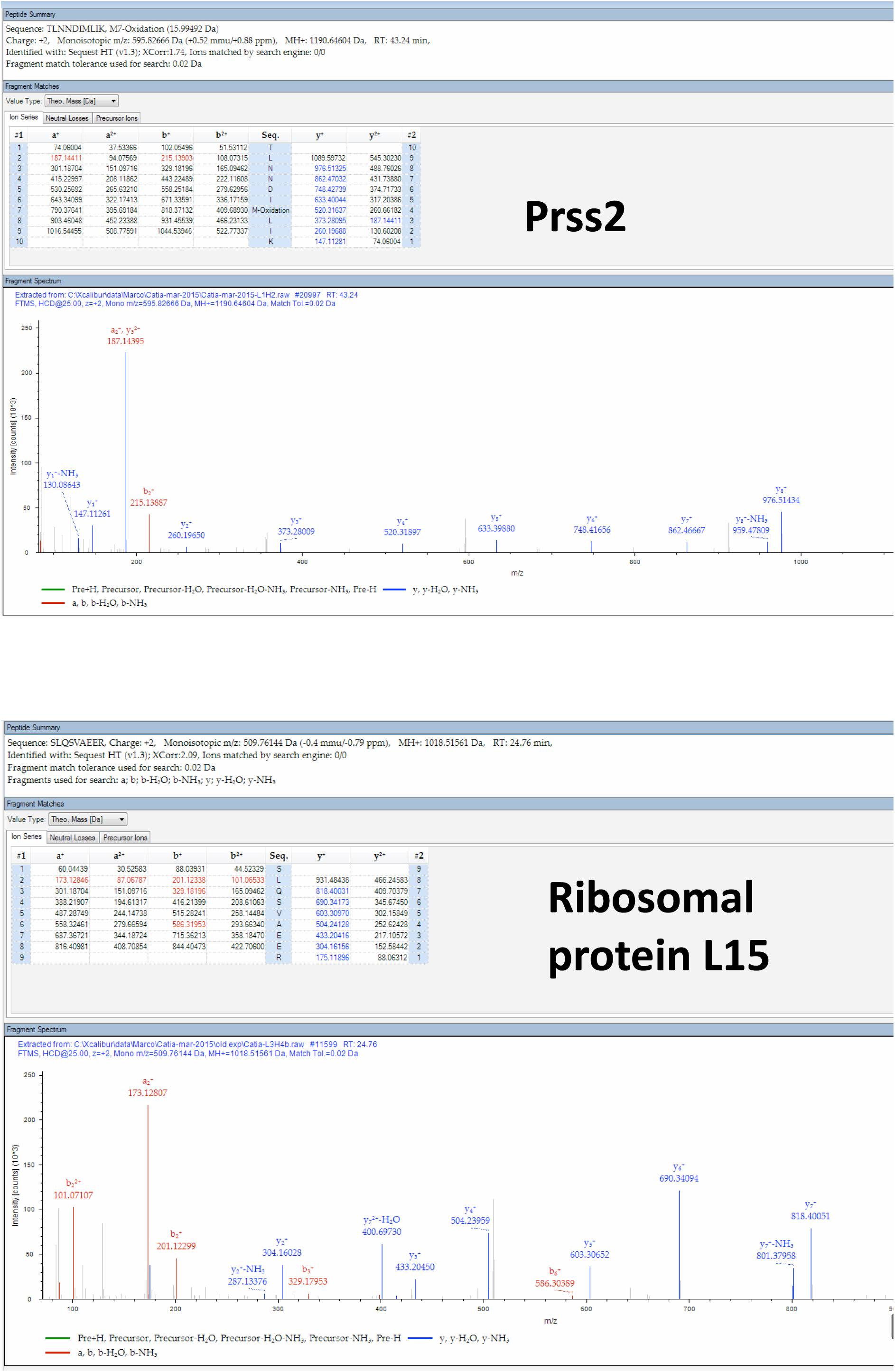
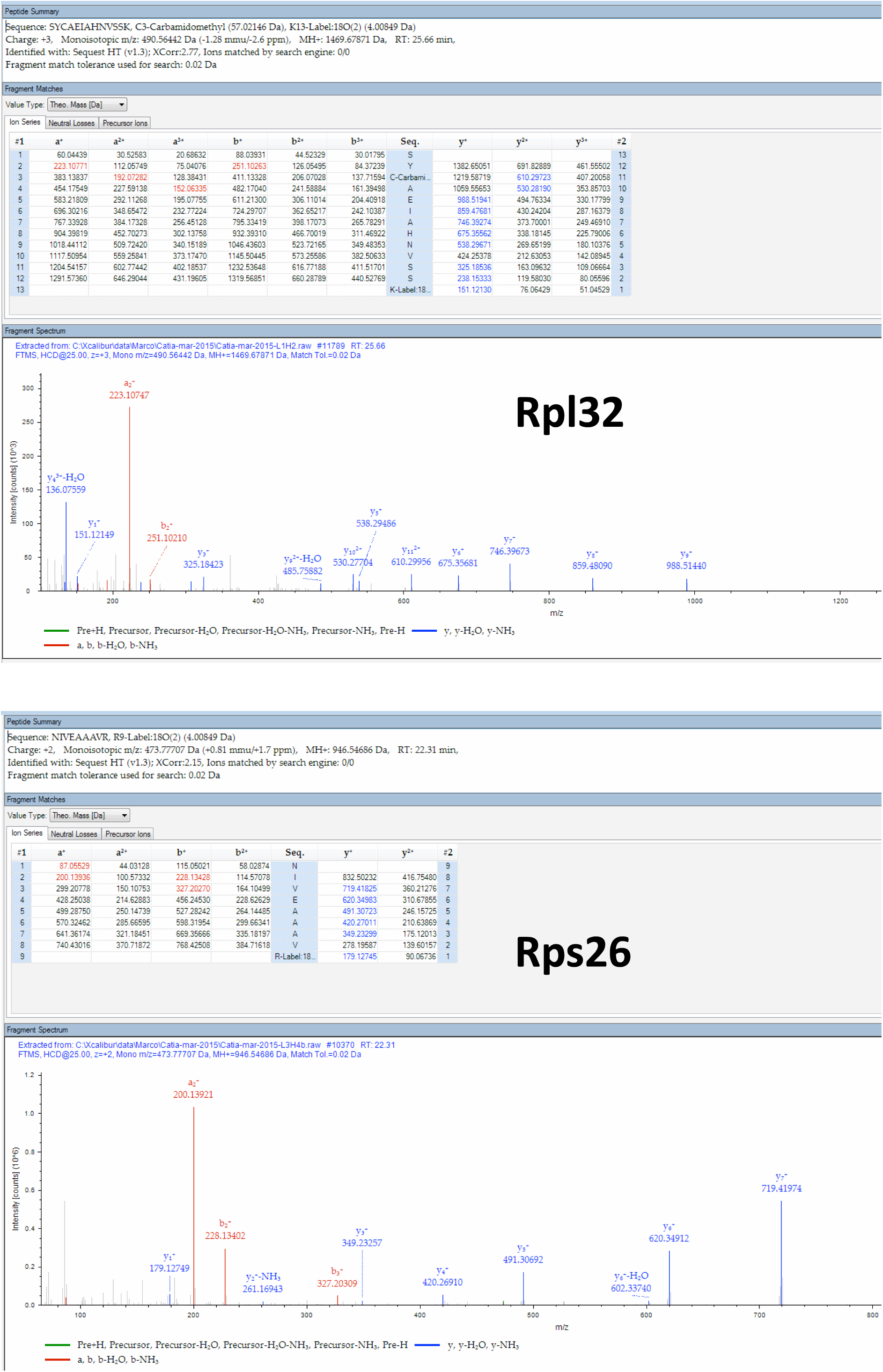
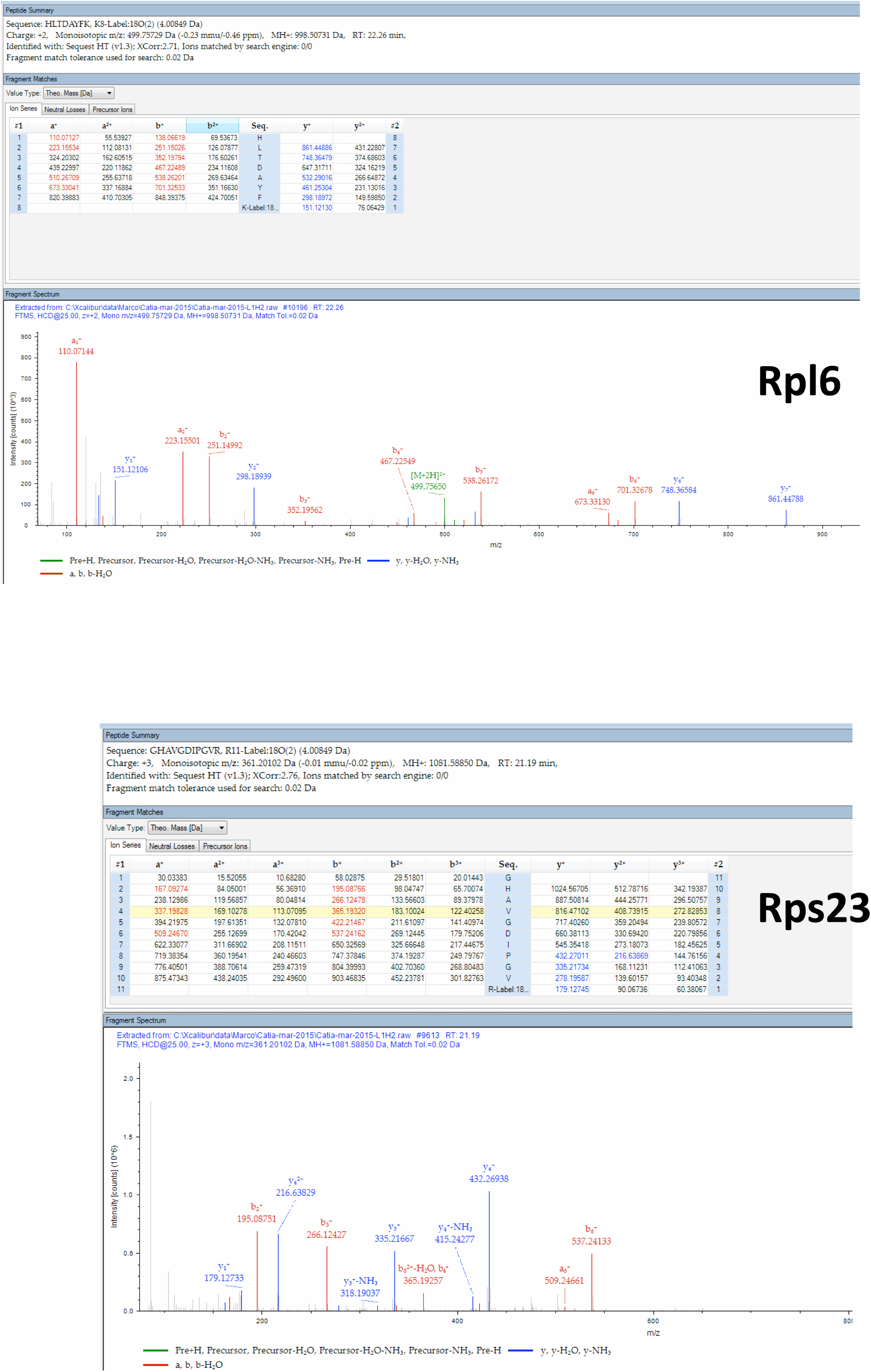
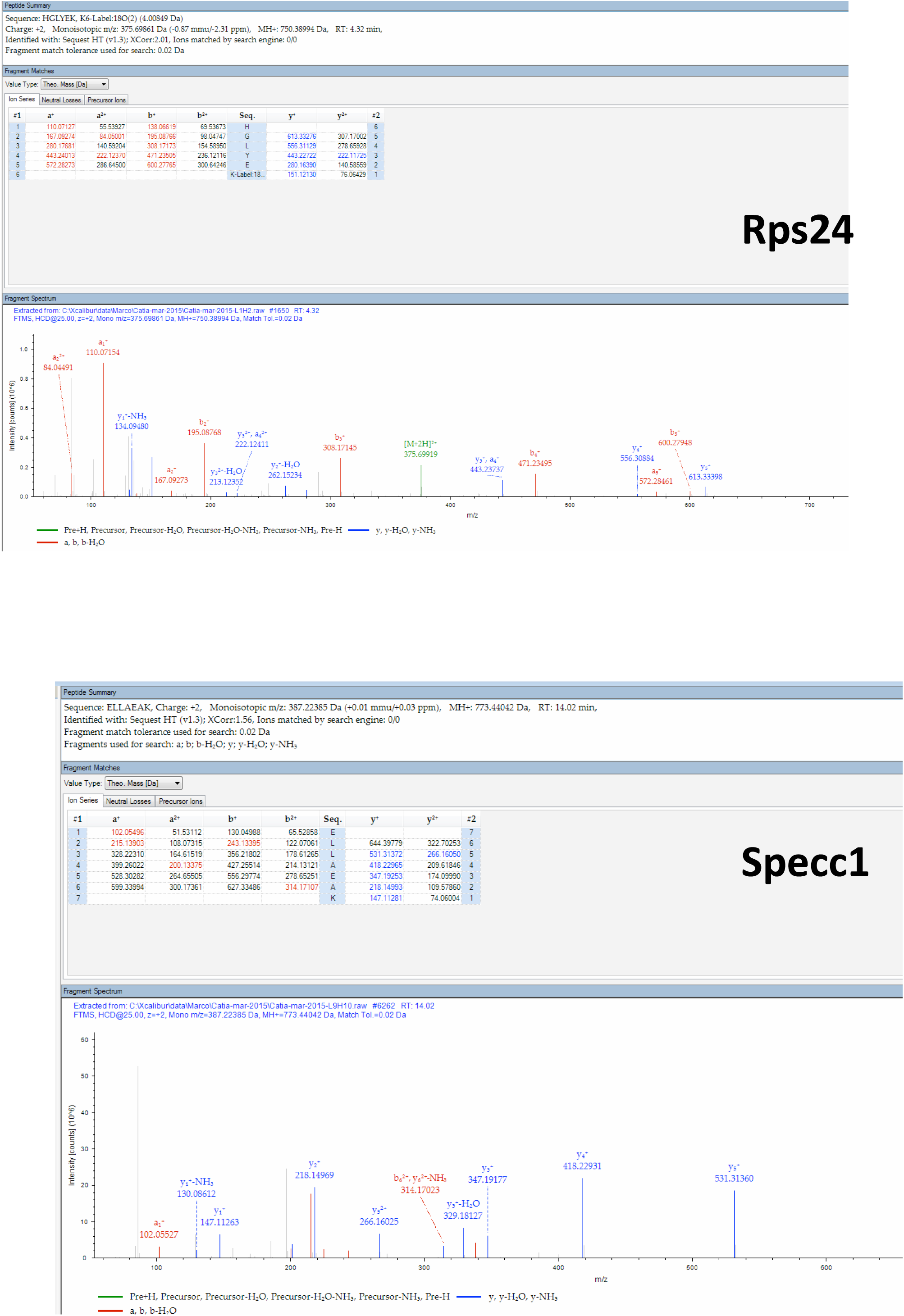
Related to Fig. 2 and S6. MS/MS spectra of protein entries identified by a single peptide match.

**Table S4:**
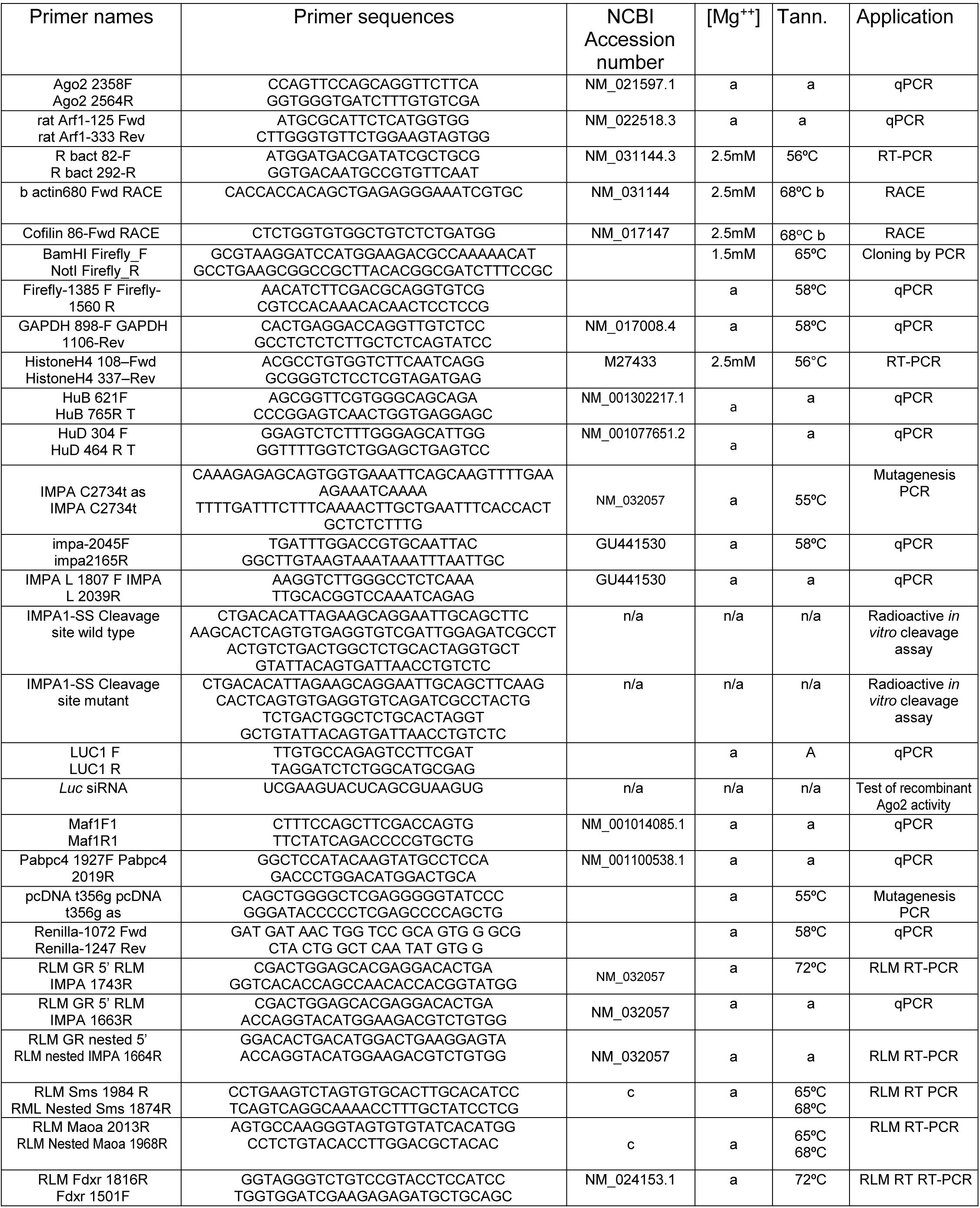
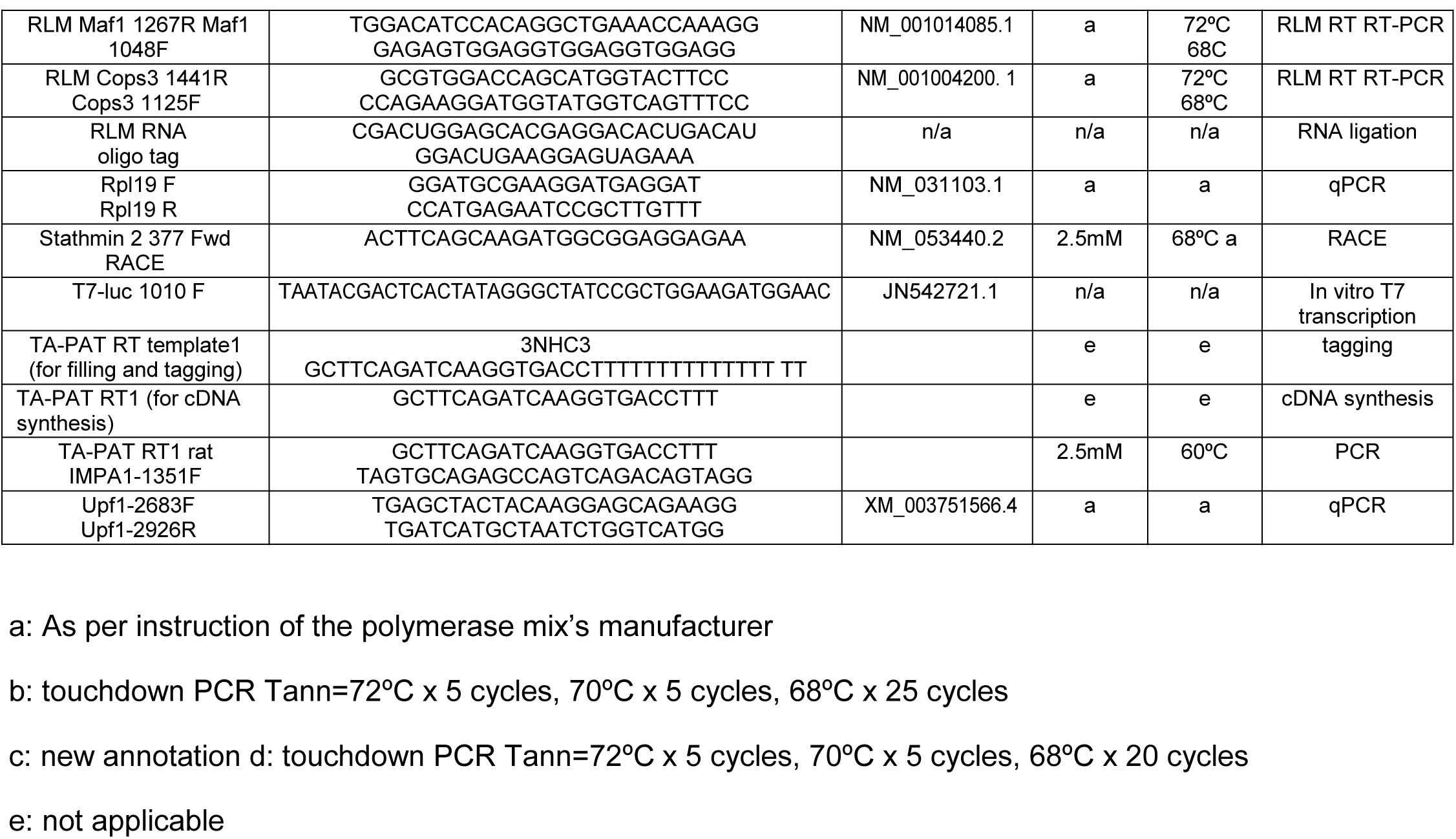
Related to Methods. Primer sequences, PCR conditions and antibodies.

**Table S5.** Related to Methods. Summary statistics for RNA-seq samples alignments.

| Compartment | Biological Replicates | n0. m1<br>[fastq] | n0. m2<br>[fastq] | n0.<br>mapped | n0.<br>alignments | n0.<br>sequences |
| --- | --- | --- | --- | --- | --- | --- |
| cell body | 1 | 9.32E+07 | 9.32E+07 | 8.27E+07 | 1.68E+08 | 7.35E+07 |
|  | 2 | 9.85E+07 | 9.85E+07 | 8.62E+07 | 1.76E+08 | 7.45E+07 |
| axons | 1 | 1.24E+08 | 1.21E+08 | 9.93E+07 | 1.85E+08 | 2.72E+07 |
|  | 2 | 1.09E+08 | 1.09E+08 | 8.08E+07 | 1.55E+08 | 2.49E+07 |

m1 left end reads m2 right end reads

## PCR and RT-PCR conditions

Initial denaturation: 94°C, 2min followed by 36 cycles [94°C, 30 sec; annealing temperature (as indicated above) 30sec; 72°C, 1min], final elongation 72°C 5min.

## Quantitative RT-PCR conditions

Enzyme activation: 50°C 2min, initial denaturation: 94°C as per manufacturer’s instruction (10 min or 2 min) followed by 40 cycles [94°C, 10sec; annealing temperature (as indicated above) 20sec; 60C°C, up to 1 min; reading], melting curve T annealing to 100°C.

### DNA oligos used for IVT of the probes used in the in vitro cleavage assay

#### Wild-type

CTGACACATTAGAAGCAGGAATTGCAGCTTCAAGCACTCAGTGTGAGGTGTCGATTGGAGATCGCCTACTGTCTGACTGGCTCTGCACTAGGTGCTGTATTACAGTGATTAA<u>CCTGTCTC</u>

#### Δcleavage site

CTGACACATTAGAAGCAGGAATTGCAGCTTCAAGCACTCAGTGTGAGGTGTCAGATCGCCTACTGTCTGACTGGCTCTGCACTAGGTGCTGTATTACAGTGATTAA<u>CCTGTCTC</u>

#### Δstem

TCAGTGTGAGGTGTCGATTGGAGATCGCCTACTGTCTGACTGGCTCTGCACTAGGTGCTGTATTACAGTGATTAATAAAACACGAAGGCCCTCTGCACAGGGAGAGCCTGCT<u>CCTGTCTC</u>

### Mutant loop

CTGACACATTAGAAGCAGGAATTGCAGCTTCAAGCACTCAGTGTGAGGTGTAAATTGGAGATCGCCTACTGTCTGACTGGCTCTGCACTAGGTGCTGTATTACAGTGATTAA<u>CCTGTCTC</u>

Underlined sequence indicates linker to anneal to T7 promoter primer according to mirVana probe kit (Thermo) instructions.

**Antibodies and dilutions**
| Antibody name | Source and catalogue number | Application and working dilution |
| --- | --- | --- |
| Anti-Ago2 | Abcam, ab186733 | WB: 1:1000 |
| Anti-GFP | Abcam, ab6556 | WB: 1:5000 |
| Rb HA | CST, 3724 | WB: 1:1000 |
| HuD | Santa Cruz, Sc-28299 | co-IP 2μg/IP<br>RIP 5μg/IP<br>WB: 1:2500 |
| Anti-neurofilament | Abcam, ab4680 | IF: 1:3000 |
| Pabpc4 | Novus Biological, NB100-74594 | WB: 1:1000 |
| PI3 kinase p85 | Upstate, 06-497 | WB: 1:2000 |
| Hsp90 | Santa cruz, sc1055 | WB: 1:1000 |
| IMPA 1 | Abcam, ab184165 | WB: 1:10000 |
| anti-αtubulin | Sigma, T9026 | IF: 1:1000 WB:<br>1:10000 |
| Anti-Upf1 | Millipore, 07-1014 | RIP 5μg/IP<br>WB: 1:2000<br>Co-IP 2μg/IP |

Applications key:
Co-IP= co-immunoprecipitation
IF=immunostaining
RIP= RNA immunoprecipitation
WB= western blotting

